# Neuro-computational mechanisms of action-outcome learning under moral conflict

**DOI:** 10.1101/2020.06.10.143891

**Authors:** L. Fornari, K. Ioumpa, A. D. Nostro, N. J. Evans, L. De Angelis, R. Paracampo, S. Gallo, M. Spezio, C. Keysers, V. Gazzola

## Abstract

Predicting how actions result in conflicting outcomes for self and others is essential for social functioning. We tested whether Reinforcement Learning Theory captures how participants learn to choose between symbols that define a moral conflict between financial self-gain and other-pain. We tested whether choices are better explained by model-free learning (decisions based on combined historical values of past outcomes), or model-based learning (decisions based on the current value of separately expected outcomes) by including trials in which participants know that either self-gain or other-pain will not be delivered. Some participants favored options benefiting themselves, others, preventing other-pain. When removing the favored outcome, participants instantly altered their choices, suggesting model-based learning. Computational modelling confirmed choices were best described by model-based learning in which participants track expected values of self-gain and other-pain separately, with an individual valuation parameter capturing their relative weight. This valuation parameter predicted costly helping in an independent task. The expectations of self-gain and other-pain were also biased: the favoured outcome was associated with more differentiated symbol-outcome probability reports than the less favoured outcome. FMRI helped localize this bias: signals in the pain-observation network covaried with pain prediction errors without linear dependency on individual preferences, while the ventromedial prefrontal cortex contained separable signals covarying with pain prediction errors in ways that did and did not reflected individual preferences.

## Introduction

We often have to learn that certain actions lead to favorable outcomes for us, but harm others, while alternative actions are less favorable for us but avoid or mitigate harms to others (1). Much is already known about the brain structures involved in making moral choices when the relevant action-outcome contingencies are well known (2–9), but how we *learn* these contingencies remains poorly understood, especially in situations pitting gains to self against losses for others.

Reinforcement learning theory (RLT) has successfully described how individuals learn to benefit themselves (10, 11) and most recently, how they learn to benefit others (12–15). At the core of reinforcement learning is the notion that we update expected values (EV) of actions via prediction errors (PE) – the differences between actual outcomes and expected values represented in mind.

Ambiguity in morally relevant action-outcome associations raises specific questions with regard to RLT, especially if outcomes for self and others conflict. If actions benefit the self and harm others, are these conflicting outcomes combined into a common valuational representation as in model-free/habit learning; or do we track separate expectations for benefits to the self and harm to others, as in model-based/goal-directed learning (16)? In addition, people differ in how they represent benefits and harms to self (17), and in whether they prefer to maximize benefits for the self vs. minimizing harms to others (3, 4, 6). How can such differences be computationally represented using RLT? Would people emphasizing one aspect, such as the harm to others, already show increased prediction errors and expected value signals when confronted with this outcome type, or are expectations tracked independently of one’s preferences, such that preferences only play out when decisions are being taken?

To address these questions, we performed two experiments: an Online experiment with 79 and an fMRI experiment with 27 participants (Table 1). In the core task common to both experiments, participants learned morally relevant action-outcome associations over different blocks of 10 trials each (Figure 1A, C; Table 1). In each block, participants saw two new symbols and faced a conflictual situation (Conflict condition): one symbol led to high monetary gains for the self 80% of the time, and to a painful but tolerable shock to the hand of a confederate with the same probability. We refer to this symbol as ‘lucrative’, since it was associated with higher monetary outcomes. The other symbol led to low monetary gains for the self 80% of the time, and to non-painful shocks to the confederate with the same probability. We refer to this symbol as ‘pain-reducing’. Importantly, to partially de-correlate representations of shock and money, the probabilities of high and low monetary reward and pain and no-pain to others were drawn independently. At the beginning of each block, participants did not know the associations between symbols and outcomes. Choosing which symbol best satisfies the moral values that participants act upon in the task thus involves learning to predict the outcomes associated with each symbol.

**Figure 1.**
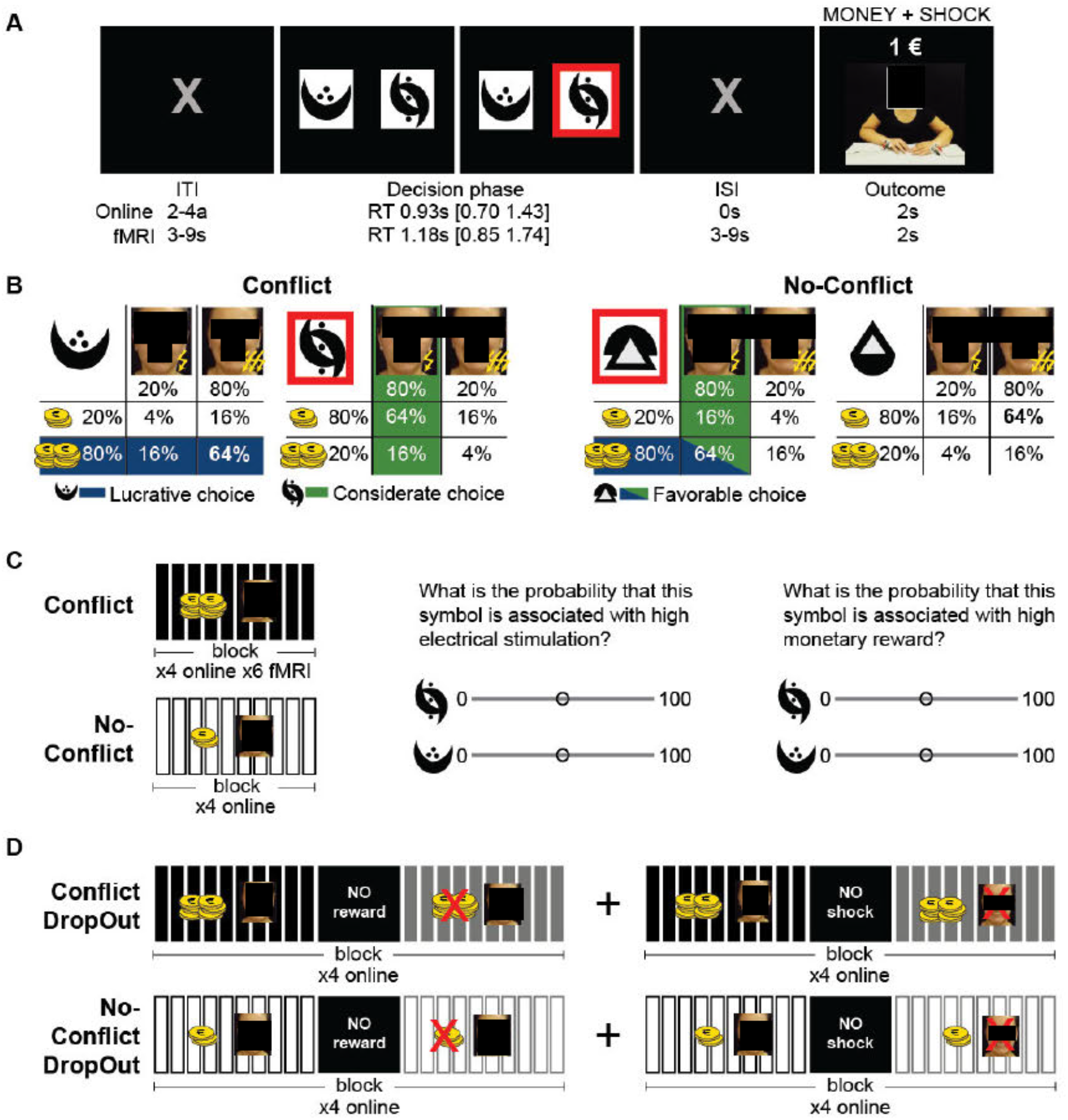
Learning Task. **(A)** Trial structure. Red outline indicates an exemplary choice for this trial. The picture showing the confederate’s response for the outcome is a still frame captured from one of the videos used in the online experiment and in which the confederate received a painful shock (see Supplementary Materials § 1 for details of the videos). Inter-stimulus and inter-trial intervals were adapted to the fMRI and online situations, and are indicated separately below each relevant instance of the trial. Reaction times (RT) are indicated for the common Conflict task. **(B)** Probability table associated with each symbol of a pair for the Conflict and No-Conflict conditions. Red outline indicates the symbol chosen in (a). Only the Conflict condition was presented in the fMRI study. **(C)** Basic block structure. Each rectangle represents a single trial. The money and face on top indicate that both outcomes were shown for that block. x4 or x6= number of block repetitions for each experiment. The questions at the end of each block were presented consecutively and separately for each symbol. **(D)** Block structure for the DropOut condition (only in the online experiment). Four blocks in which the money and four in which the shock to the other were removed were presented in a randomized order. Black indicates trials in which both money and shock to others are presented, gray those in which either money or shock is removed.

**Table 1.**
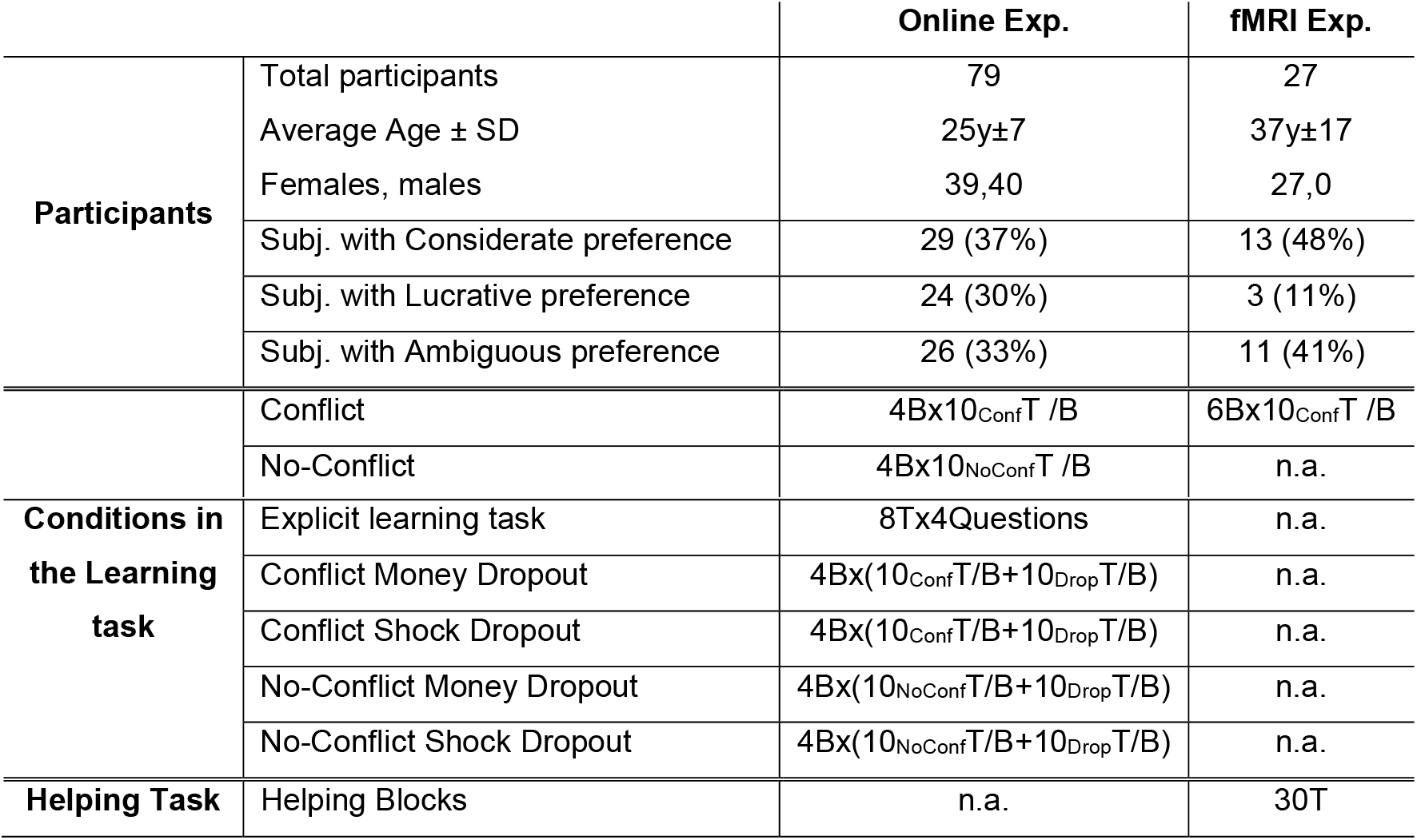
Participants and Task-conditions overview. For each experiment, the table reports the total number of participants included in the behavioural analysis, and how many of those fell within the Considerate, Lucrative and Ambiguous preference pattern as calculated by the Binomial distribution. Additionally, the table reports the number of repetition of each condition included in that particular experiment. T=trial; B=block; Conf=Conflict; NoConf=NoConflict; Drop=Dropout. fMRI data only included the right handed participants, 25 (35y±15SD; 25f), while behavioural data were collected for the 27 indicated in the table. Note: although age differed between the fMRI and Online experiment (Mann-Whitney test, BF_10_=8.022, p<0.001), the choices did not differ significantly (Kendall’s tau(age, %Considerate choices)=0.113, BF_10_=0.543, p=0.096).

In the online experiment, participants performed a number of additional tasks. First, to explore whether participants learned the symbol-outcome association probabilities for both the self-money and other-shocks, even if their preferences may emphasize one, we added blocks in which we crossed the association to remove the conflict. In this No-Conflict condition (Figure 1B), the symbol that led to high money in 80% of cases also led to low shocks in 80% of the cases, and the symbol leading to low money in 80% of the cases also led to high shocks in 80% of the cases. After each of these blocks we then explicitly asked participants to report the learned associations (Figure 1C). Only if participants learn symbol-outcome association for the less preferred outcome, will they report different probabilities under the Conflict and No-Conflict for this outcome. Second, to distinguish model-based from model-free learning, we adapted the devaluation approach pioneered in animals (16). In a classic example of this approach (18), animals learned to press a lever for an attractive food A. All rats then underwent taste aversion learning, but for some, food A, the one they worked for, was paired with a lithium-chloride injection, for others the injection was paired with a different food B. When both groups could later press the lever again, only those for whom food A had been devalued reduced their usage of the lever. This showed their behavior was model-based: driven by an understanding that the lever didn’t simply lead to rewarding states, but to food A - a food that had lost its value and wasn’t worth working for anymore (18). Here, we therefore added blocks (DropOut condition, Figure 1D) in which after 10 trials to learn the symbol-outcome associations, we informed participants that the self-money or the other-shocks would not be delivered on the following trials. We then examined the choice on the 11^th^ trial, before participants witnessed the modified outcome. If participants use a model-based strategy, we expect them to show different choices depending on which outcome is removed, with choices expected to change substantially from the 10^th^ to the 11^th^ trial, if the outcome they weigh more is removed. If participants use a model-free strategy, as has recently been suggested using a two-stage decision-task(15), we expect them to continue choosing their previously favoured option. The comparison of participant’s choices at the 11^th^ trials between the Conflict and NoConflict conditions, further informs us on their ability to form separate representations for money and shocks.

For the participants in the fMRI experiment, beside the core learning task, we also performed a different costly helping task previously developed in our group (4), to test whether parameters we extract from our conflictual learning task can predict choices in a different costly helping task not requiring learning (Supplementary Material §3).

After describing participants’ behavior, we used Hierarchical Bayesian Model comparisons to test which alternative computational formulations of the RLT models (Figure 2) better describe participant’s choices, and gain insights into how people combine the two outcomes in their morally relevant learning experience. Except in our random choice model (M0), in all the learning models we compare, shock and money are additively combined using an individual weighting factor *wf* ranging from 0 to 1. This *wf* captures the value of the monetary outcome for self relative to the value of shock to the other and is not unlike the salience alpha in the Rescorla-Wagner Learning Rule (19, 20). A *wf* closer to zero would characterize a participant preferring to minimize the harm to the other person, a *wf* closer to one corresponds to more lucrative choices. In particular, we investigate whether choices are better predicted (i) by an RLT model that combines money and shock as soon as outcomes are revealed (M1) or (ii) models that keep separate representations for the two types of outcomes (M2). M1 instantiates model-free learning in which participant’s decisions are based on the history of past reward values, without representation of the nature of outcomes, while M2 instantiates model-based learning with separable representations of the expected outcomes for self and others. For M2, we further compared a variant that scales outcomes based on personal preferences for money vs. shock (M2Out) vs. a variant that initially tracks expectations independently of personal preferences regarding outcomes, but that introduces weights at the decision phase of the task (M2Dec). We expect all but M0 to perform comparably well during the conflict phase of our experiment. We however expect that if participants use model-based learning under conflict, M2 should outperform M1 during the 11^th^ trial when one outcome is removed. We had no specific predictions about whether M2Dec or M2Out would predict choices at the 11^th^ trial more accurately.

**Figure 2.**
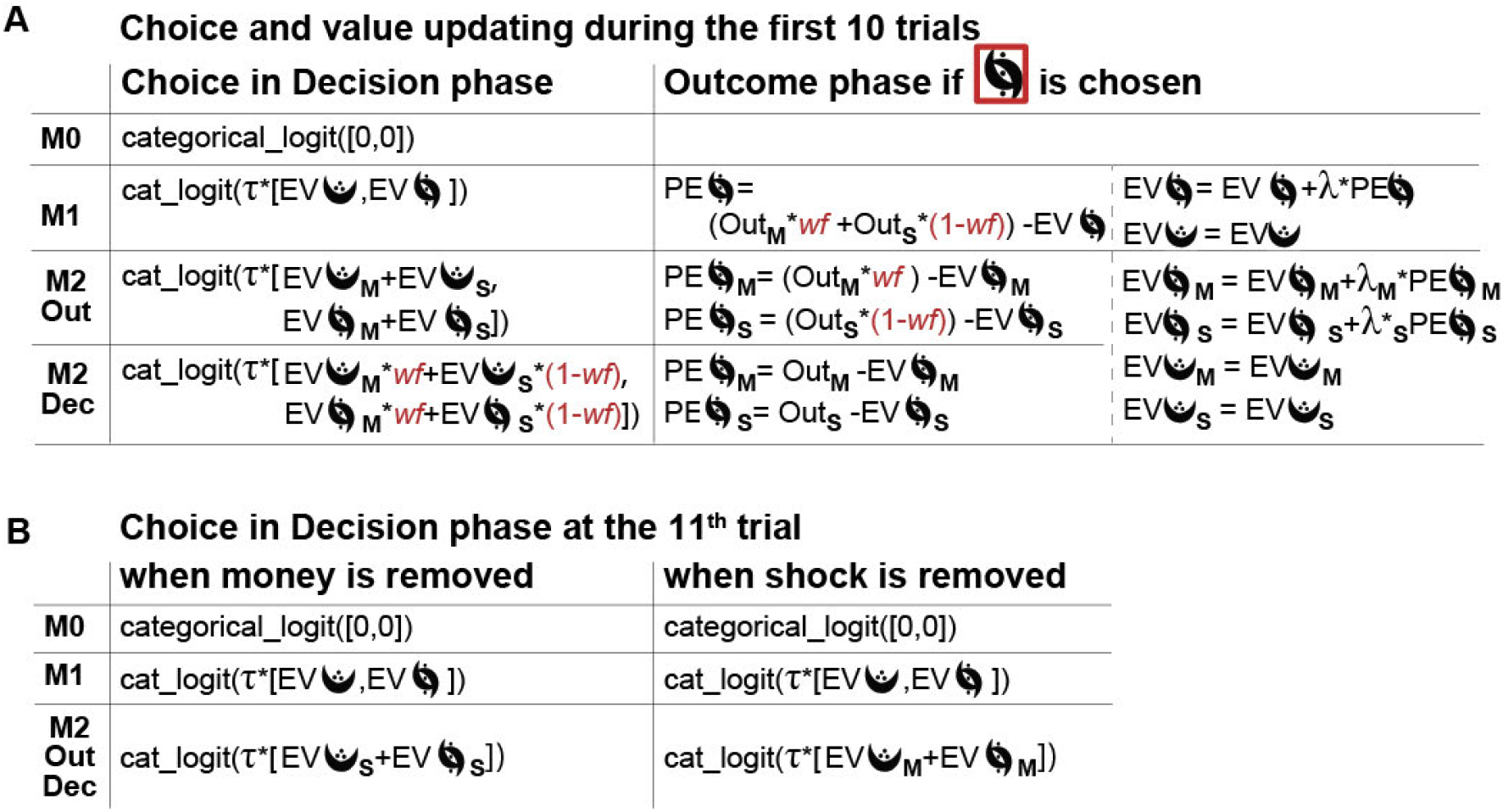
Tested RLT Model. **(A)** Model formalization for the symbol chosen in Figure 1a. In all our models, decisions are based on a categorical logit function (aka softmax), contrasting the expected value across the alternatives with inverse temperature τ□[0,5] such that for a given difference in expected value, participants with higher τ choose the symbol with higher expected value more often. The learning rate λ□[0,1] captures the speed at which expected values are updated, with higher λ values indicating that expected values are more strongly updated following surprising outcomes. EV=Expected Value, PE=Prediction Error, Out=Outcome. Subscript M=money, Subscript S=shocks. Outcomes are coded based on their value, such that high-shock OutS=-1, low-shock OutS=+1, high-Money OutM=+1, low-money OutM=-1. Trials in which one outcome is withheld during dropout is coded as Out=0. Note that outcomes are set to the same values (+1 and −1) for money and shock because we chose the amount of money at stake to be close to the indifference point (see Supplementary Materials §2). **(B)** Model formalization for the symbol chosen in Figure 1a during the decision phase at the 11^th^ trial for the DropOut condition.

Shocks were made visible to the participants through pre-recorded videos showing facial expressions from the confederate. In the fMRI experiment participants believed these recordings to be part of a real-time video-feed, while in the Online experiment they were aware that the videos had been pre-recorded, but believed that after the task we would select 15 trials and administer the corresponding shocks to another participant in the lab. We used this video-feedback, instead of the symbolic feedback more often used in neuroeconomic paradigms, to explore the neural systems activated in situations where the consequences of our actions are only available from the facial expressions of the people around us, and to remain closer in design to the paradigms used in nonhuman animal models (21).

Finally, we used the neuroimaging data to inform our understanding of how our participants update values in our tasks. Given substantial literature on how individuals learn to maximize gains for the self (22, 23), we focus on how the brain updates values based on shocks to others. In particular, given the conceptual difference between M2Out and M2Dec, we ask whether we can distinguish and localize areas in which BOLD signals covary with prediction errors in ways that do vs. do not depend on *wf*. More specifically, given an extensive literature that has outlined a network of brain regions recruited while witnessing painful vs non-painful facial expressions (24–27), and the availability of a multivariate signature to quantify activity in this network (28), we first ask whether activity in this network covaries with prediction errors for shocks, and if so, whether it does so in ways that do or do not depend linearly on *wf*. Second, we perform a more explorative analysis to localize where in the brain the BOLD signal covaries with pain prediction errors in ways that do or do not significantly depend linearly on *wf*. Recent reviews identified that the ventromedial prefrontal cortex (vmPFC) has BOLD signals that covary positively with the current value of multiple outcomes, in particular for chosen options (22, 29), and outcomes for others appear to also be encoded in this region (30, 31). Interestingly, when switching between making choices to benefit the self and others, the most ventral medial prefrontal cortex had BOLD signals covarying with the value that was at the focus of a particular trial while a more dorsal part covaried with the irrelevant value (32). In our task we may thus expect the ventral mPFC to have signals that covary with prediction errors for shocks in ways that depend on *wf* (and at thus at the focus of a particular trial) while more dorsal mPFC regions may also represent prediction errors for shocks in ways that do not depend on (which may become relevant in the future if contingencies change).

## Results

### Participants vary in their preference but most favour one symbol in the first 10 trials of a block

Figure 3A-B shows participants’ choices in the Conflict condition (see also Table 1). Using a binomial distribution, with 60 trials for the fMRI experiment (10 trials x 6 blocks), and 120 trials for the Online experiment (10 trials x 12 blocks), we found that in the fMRI experiment about half the participants chose the pain-reducing option above chance, about half were within what would be expected without preference, and just 3 chose the lucrative option above chance. For shorthand, we refer to the choice patterns in these subgroups, divided by the binomial probability, as ‘considerate’, ‘ambiguous’ or ‘lucrative’ preferences. In the Online experiment, participants were evenly subdivided across these three preference subgroups. Averaging the learning curves of participants showing opposite preferences would occlude learning, and Figure 3B thus visualizes the average choices for each group separately. The groups with lucrative and considerate choice preferences show typical learning curves, starting at chance (50% preference for the pain reducing option) on the first trial of each block and then gravitating towards the pain reducing (for considerate) or lucrative (for lucrative preferences) option, with the last four trials showing relatively stable preferences around 80%, as would be expected for an RLT with 80/20 probability. The curve of the group with ambiguous preference, consistently remains around 50% showing no clear learning curve (Figure 3B, orange and yellow). Finally, participant’s average choices did not seem to be influenced by how much participants believed in the shock delivery (Supplementary Materials §4), and were in agreement with participant’s reported motivation (Supplementary Materials §5).

**Figure 3.**
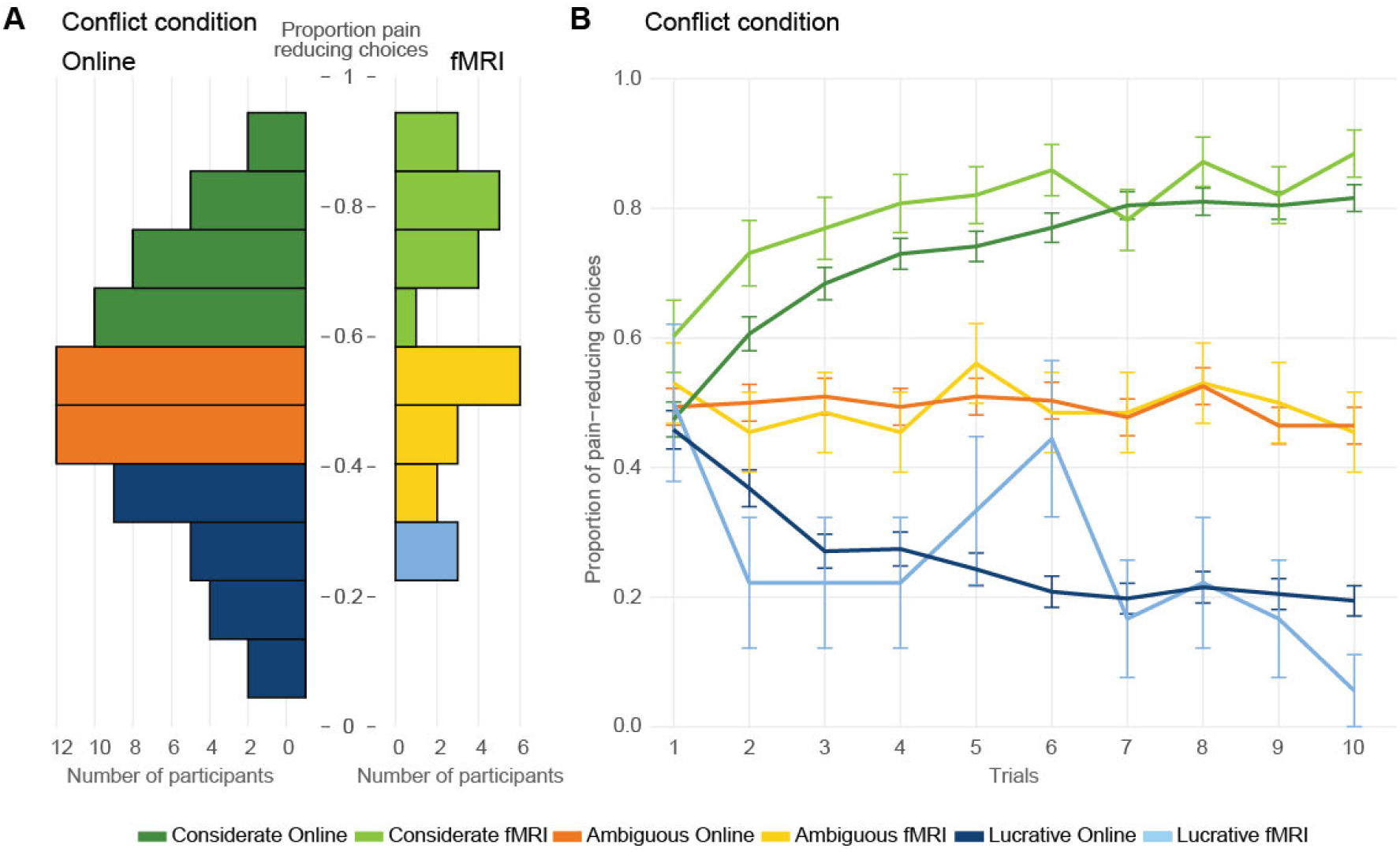
Choice allocation in the Conflict condition. **(A)** Histogram showing the proportion of pain-reducing choices per participant over the first 10 trials of all available blocks of the online (left, 12 blocks of 10 trials) or fMRI (right, 6 blocks of 10 trials) experiment. For blocks with DropOut, only the first 10 trials, before DropOut occurs, are included (black rectangles in Figure 1D). Participants with above chance proportion of pain-reducing choices according to a binomial (*p*<0.05, considerate preference) are shown in dark (online) or light (fMRI) green, participant with below chance (*p*<0.05, lucrative preference) are shown in dark (online) or light (fMRI) blue, and participants within chance level (*p*>0.05, ambiguous preference) are shown in orange (online) or yellow (fMRI). **(B)** Choices as a function of trial number within a block, averaged over blocks, separately for the three preference subgroups, with error bars representing the s.e.m. over participants. Colour-code as in (A). As in (A) all first 10 trials of the Conflict conditions are included, independently of whether they belong to a DropOut or No-DropOut block. For the comparison of choice allocations across conditions see Figure 3_figure supplement 1.

### Participants have explicit and separable access to the symbol-outcome associations for money and shock that are biased by preference

To probe participants’ explicit learning, in the Online experiment we asked them to report how likely each symbol was associated with high-shock and high-money. Overall, participants tended to report higher probabilities for symbols with higher probability, also capturing the difference between the Conflict and NoConflict condition (comparing the *N* and U shapes in Figure 4A). Because choices are thought to be driven by the *difference* in expected value across options, we summarized explicit reports as the difference in reported probability for the favored minus the non-favoured symbol, with larger reported differences in the correct direction providing more evidence for (explicit) learning (Figure 4B). We observed that the reported differences in probability across the symbols was different from zero, and in the expected direction in all cases (All BF_10_>3), showing that as a group, participants with considerate and lucrative preferences also learned something about the association with outcomes that they appear to be less interested in. Second, participants show evidence of noticing the Conflict vs No-Conflict difference, which they would not need to do if they focused exclusively on maximizing money or minimizing shocks: participants with considerate preference (green) reported lower association with high-shock for their preferred (shock-reducing) symbol in the conflict and no-conflict block (both 3^rd^ and 4^th^ violin below zero), and they reported that their favored symbol had lower high-money probability in the Conflict (1^st^ violin) and higher high-money probability in the No-Conflict (2^nd^ violin) than their non-preferred symbol. A similar pattern is true for participants with lucrative preference (blue): they reported that their favorite (money-maximizing) symbol had a higher high-money probability than their non-favored symbol after both Conflict and No-Conflict blocks, and that this symbol was more likely in Conflict and less likely in No-Conflict blocks to lead to high-shock than their non-preferred symbol. Also participants with ambiguous preference could report above chance differences in probability for the two symbols in the correct direction in all cases (Figure 4C).

**Figure 4.**
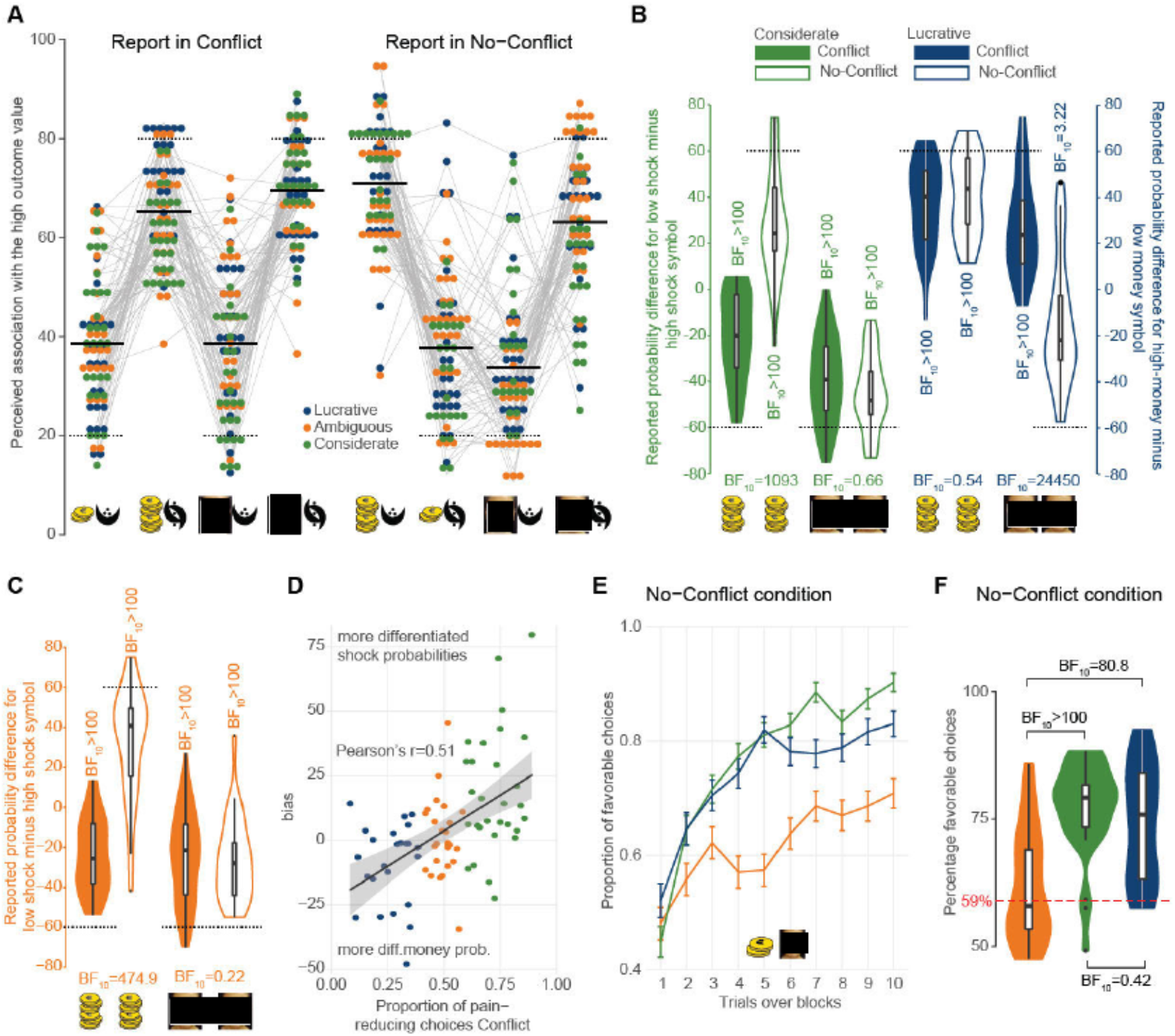
Participant reports and No-Conflict choices. **(A)** Participants’ reported perceived high-outcome probability for both symbols (see Fig. 1C) from the Conflict (left) and No-Conflict (right) conditions. The x-axis specifies which symbol and question were asked: short/tall pile of money=reported high-money probability for the symbol actually associated with low/high-money; painless/painfiil face=reported high-shock probability for the symbol associated with low/high-shock. As a group, most participants correctly assigned higher probabilities to symbols that had higher probability, and reported a different pattern of probabilities after Conflict and NoConflict blocks. Thick black lines = average of reported probabilities: dotted black lines = programmed/expected probability. **(B)** Difference between reported probabilities for the preferred minus non-preferred symbol separately for the Considerate (green) and Lucrative (blue) preference group and for the Conflict (filled violins) and No-Conflict (empty violins) condition. The symbols on the x-axis specify whether the question was to estimate the probability of leading to high-money (coins) or high-shock (face). Violin plots report median and quartiles. The BF above a violin represents the result of a Wilcoxon signed rank test of the differences vs. zero; the BF below a pair of violins represents the result of a Wilcoxon signed rank test comparing the conflict and no-conflict conditions. Figure 4_figure supplement 1 guides the reader through the computations presented in this panel. Dotted black lines = programmed probability. **(C)** Same as in (e) for the ambiguous group. **(D)** Correlation between the participant’s report bias and the proportion of pain-reducing choices during the conflict task, with Bias= (rpS(high shock) - rpS(low shock)) - ((rpM(high money) - rpM(low money)), where rpS/rpM stand for reported probability of high-Shock/high-Money outcome, and highshock, lowshock. highmoney. lowmoney refer to the symbols based on their actually most likely outcome. **(E)** Choice in the No-Conflict condition. Note that the classification in terms of preference is still based on the Conflict choices. **(F)** Proportion of favorable choices (i.e. choosing the symbol most likely to lead to high-money and low-shock) separately for the three groups.

Interestingly, the magnitude of explicitly reported probability difference was biased in favour of the outcome participants seem to weigh more strongly. When separating participants in preference-groups, we see that those choosing the pain-reducing option above chance show a stronger difference, that was also closer to the actual difference, in reported probability for the shock than money outcomes, and those choosing the lucrative option show a stronger difference, that was also closer to the actual difference, for the money than the shock outcomes (difference between the filled green violins in Figure 4B for the considerate preference= 18.2±4.45s.e.m, W=377.5, BF_10_=577.22, p<0.001 and between the filled blue violins for the lucrative=9.2±3.25s.e.m., W=233, BF_10_=7.67, p<0.018). Because the thresholds for grouping participants in these three preference groups is somewhat arbitrary, we also examined whether over all participants, the preference (calculated as proportion of pain-reducing choices) was predictive of their bias, and this was the case (Figure 4D, Pearson’s r= 0.51; t_(77)_=5.2, p= 1.6 x 10^-6^).

That ambiguous preference participants could still report above chance probability estimates (Wilcoxon signed-ranked, all p<0.001, BF_10_>100; Figure 4C) suggests that their lack of above chance choice preference was not due to a total lack of symbol-outcome association learning. A lack of clear preference for one outcome over the other seems also to have played a role. Indeed, when the conflict was removed (Figure 4E orange), ambiguous preference participants do demonstrate learning, but choose the favorable option less frequently than the other two groups (Bayesian one way ANOVA, F_(2,76)_=15.6, p<0.001; BF_incl group_=8284.52, Figure 4F). At the single subject level, in No-Conflict blocks, 54% of the ambiguous preference subjects fell below the learning threshold determined using a binomial distribution (71/120 correct choices; red line in Figure 4F), while just 7% of those with considerate and 4% of those with lucrative preference were below this threshold. The ambiguous preference group thus appears to contain a mix of individuals that do not show significant learning in our task even without conflict, and individuals that can learn without conflict but fail to display clear preferences in the presence of conflict.

### DropOut trials suggest model-based learning with separate outcome representations biased by preference

Examining the choices on the critical 11^th^ DropOut trial, similar to a classic devaluation procedure, speaks in favour of a model-based learning by confirming that participants have separable representations of self-money and other-shock, and have also learned more about the outcome that drives their decisions than about the other outcome. We find that for the considerate and lucrative groups, which showed strong preferences on the 10^th^ trial, removing the quantity that guided their choice most, led to a highly significant change in their choice allocation (Figure 5A, both BF_10_>80), while removing the quantity that guided their choices less, left theft choices unchanged (both BF_10_<1/3).

**Figure 5.**
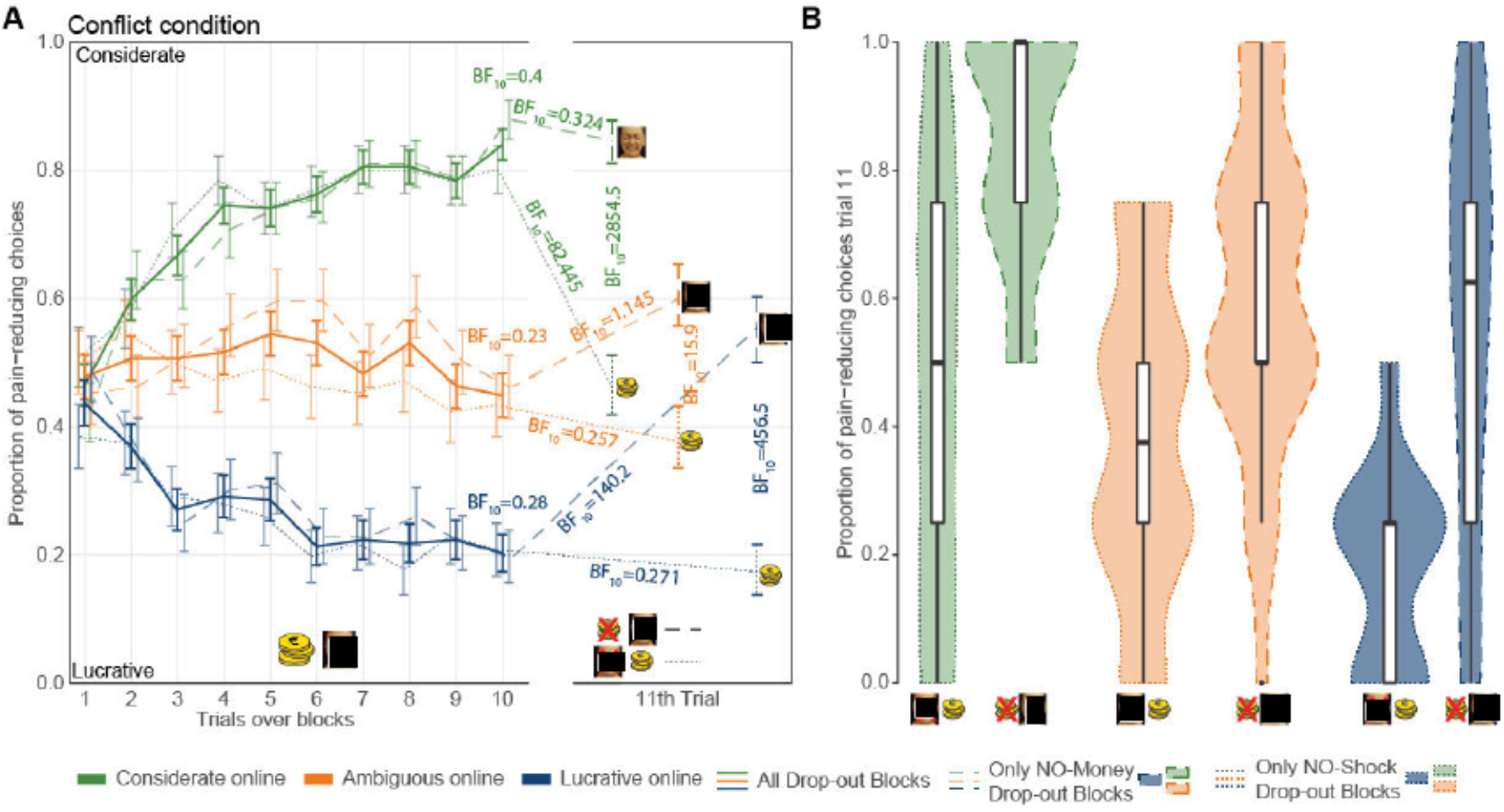
Participant’s choices at the 11^th^ trial. **(A)** Choices in the Conflict DropOut blocks averaged separately for the preference subgroups as classified using all Conflict trials. For each subgroup, the solid line represents the average of all trials, the dashed line, blocks in which money will be withheld, and the dotted lines, blocks in which shocks will be withheld. BF_10_ values shown between the 10^th^ and 11^th^ trial indicate the result of a Bayesian paired t-test (i.e. a Wilcoxon signed-rank test on the differences) comparing choices in these two trials. BF_10_ value shown above the 10^th^ or the 11^th^ trial, the result of the same test comparing choices on trials wrhere money will be or is withheld vs trials where shocks will be or are withheld. Error bars are the s.e m. across participants. **(B)** Violin plot of choices across participants on the 11^th^ trial of conflict drop-out blocks as a function of preference subgroup and outcome withheld. The boxplot within each violin signifies the 4 quartiles of the data, the black dot represents one outlier.

Dropout trials in NoConflict blocks did not lead to comparable changes in choices (Supplementary Figure 4; all 11^th^ vs 10^th^ trial BF_10_<2.1), confirming that the change in choices in the Conflict condition are not the result of a simple rule ‘reset choice when my prefered preferred outcome is removed, but not when my non-preferred outcome is removed’, but considers the probability of outcomes associated with the remaining outcome, which are the only difference between Conflict and No-Conflict blocks.

Interestingly, removing the guiding outcome in Conflict trials did not lead to a mirror symmetrical (i.e. ~80%) preference for the other symbol: the average choice allocation switched from ~80% (considerate) or ~20% (lucrative) pain-reducing choices on the 10^th^ to just below, or above, ~50%, on the 11^th^, respectively (Figure 5A). That the first choice after DropOut is not as polarized as the 10^th^ trial is in line with the bias we observed in the explicit reports (Figure 4B,D): given that the difference in reported outcome probabilities was less pronounced for the less-guiding outcome, one would expect less polarized choices in a decision based on these lesser differences in expected values. For the ambiguous group, there was no robust evidence that removing either quantities lead to a change of preference towards the better option for the remaining outcome. It is important to emphasize that the choice on trial 11 occurs before the participant is presented with the single remaining outcome, and thus probes choices exclusively based on expected values that were learned during conflict trials. Supplementary Figure 4 also shows choices on trials 12-20, but as these trials do not contain two potentially conflicting outcomes, but only one outcome, we do not expect our models to adequately represent participants’ reasoning and learning over those subsequence trials.

In summary, choices reveal substantial individual variability in preference, but most participants show evidence of some form of learning. Explicit reports and their difference across Conflict and No-Conflict blocks reveal that participants have explicit and separable representations of outcome probability differences across symbols in terms of money and shock, although these representations are more differentiated for the outcome that their choices indicate to bear more weight. Finally, devaluation performed by informing participants that their preferred outcome will no longer be delivered leads to an immediate switch away from their previously preferred option during Conflict. Together this supports the notion that choices in our conflictual conditions are dominated by model-based learning in which participants represent separable outcomes for money to self and shock to others, with the individual preferences influencing the magnitude of their respective symbol-outcome association. This would suggest that M2Out (Figure 2) might be the computational implementation of RLT learning most suitable to capture our participants’ choices in our paradigm.

### M2Out predicts participant’s choices more accurately

Devaluation, as implemented in our conflict Drop-Out blocks, is critical to distinguish model-based and model-free learning. To directly compare our neurocomputational models (Figure 2A), we therefore fitted our competing models on the first 10 trials of the Conflict DropOut blocks, and then examined (i) how well they predicted those first 10 choices using an approximation of the leave-one-out information criterium (LOOIC(33)) and (ii) how well they predicted choices on the 11^th^ trial that was not included into the model fitting, when one of the outcomes is removed. Because the 11^th^ trial was not included in the fit, we can directly quantify the likelihood of the observed choices on the 11^th^ trial under the different models to assess which model best predicts behavior under devaluation. *A priori*, we expect the four models to make very different predictions on the 11^th^ trial. M0, by design, continues to predict random choices. Because M1 does not have separable representations of money and shock, its predictions for the 11^th^ trial are simply based on the symbol with highest composite expected value (EV) based on the first 10 trials, and thus predicts that choice behavior does not change on the 11^th^ trial. Finally, because the M2 models have separate EV for money and shock, we programed the model to transform the information that participants receive *before* they perform the 11^th^ choice (i.e. one outcome will no longer be delivered), into a revised decision criterion based on the remaining expected value only, without using a *wf* in their choice, as there is no longer a conflict to resolve (Figure 2B). For M2Out and M2Dec, choice is then dictated for the 11^th^ trial by cat_logit(τ(EV_R_(symbol1),EV_R_(symbol2)), where subscript R represents the remaining quantity: M (money) or S (shock). We thus expect choices not to change much if the less preferred outcome is removed, and to change significantly, if the preferred outcome is removed.

Figure 6A illustrates the predictions of the three learning models (M1, M2Out, M2Dec) as compared with the actual choices for the critical DropOut blocks. For visualization purposes, we did not include M0 and the ambiguous preference participants in this figure, because they do not predict or show the learning behavior we are trying to capture. During the initial 10 trials in which the conflict is present, all three learning models (M1, M2Out, M2Dec) capture the general shape of the learning curve, and can accommodate individual differences in preference through the weighting factor *wf*, generating increasingly more pain-reducing choices as the trial number increases for participants with considerate preferences, and fewer pain-reducing choices for participants with lucrative preference. Including all participants (including the ambiguous participants), model comparison over the first 10 trials with the LOOIC confirms that M1, M2Out and M2Dec perform similarly well, i.e. remain within a standard error of one-another, and perform better than the random-choice model M0 (Figure 6B, and Supplementary Figure 5 for results that exclude the ambiguous and non-believers groups). Note that the information criterion (IC) scale captures how much information is lost when comparing model predictions with actual choices, and smaller IC thus characterize models that better describe behavior. A similar pattern is true for the choices in the fMRI experiment (Supplementary Figure 5c). Next, we used the 11^th^ trial to arbitrate across M1, M2Out and M2Dec. As we saw in Figure 5, participants’ actually responded to devaluation in ways that depended on their preference. Lucrative preference participants kept their choices unchanged when shocks were removed, but switched their preference to just above 50% pain reducing choices on average when money was removed. The converse was true for Considerate preference participants. When money is removed and only shocks remain (Figure 6B, gray background), all learning models correctly predict that considerate preference participants do not change their choices, but only M2Out (purple, and marked by the black arrow-head) accurately predicts that participants with lucrative preference change their preference to just above 50%. In contrast, M1 (gray) fails to predict any change for those lucrative participants, and M2Dec (light blue) overestimates their change. When shocks are removed and only money remains (Figure 6B yellow background), all learning models correctly predict a lack of change for the lucrative preference participants, but again, only M2Out correctly predict the magnitude of change in the considerate participants, that now choose the pain-reducing symbol just under 50% of cases on average, while M1 fails to predict any change and M2Dec overestimates the change. When considering all participants’ choices (including the ambiguous), Figure 6C shows that M2Out consistently predicts the choices on the 11^th^ trial better than any of the other models. M2Out also outperformed the other models when excluding ambiguous participants or those that signaled some degree of scepticism about whether someone would really be administered shocks (Supplementary Figure 5).

**Figure 6.**
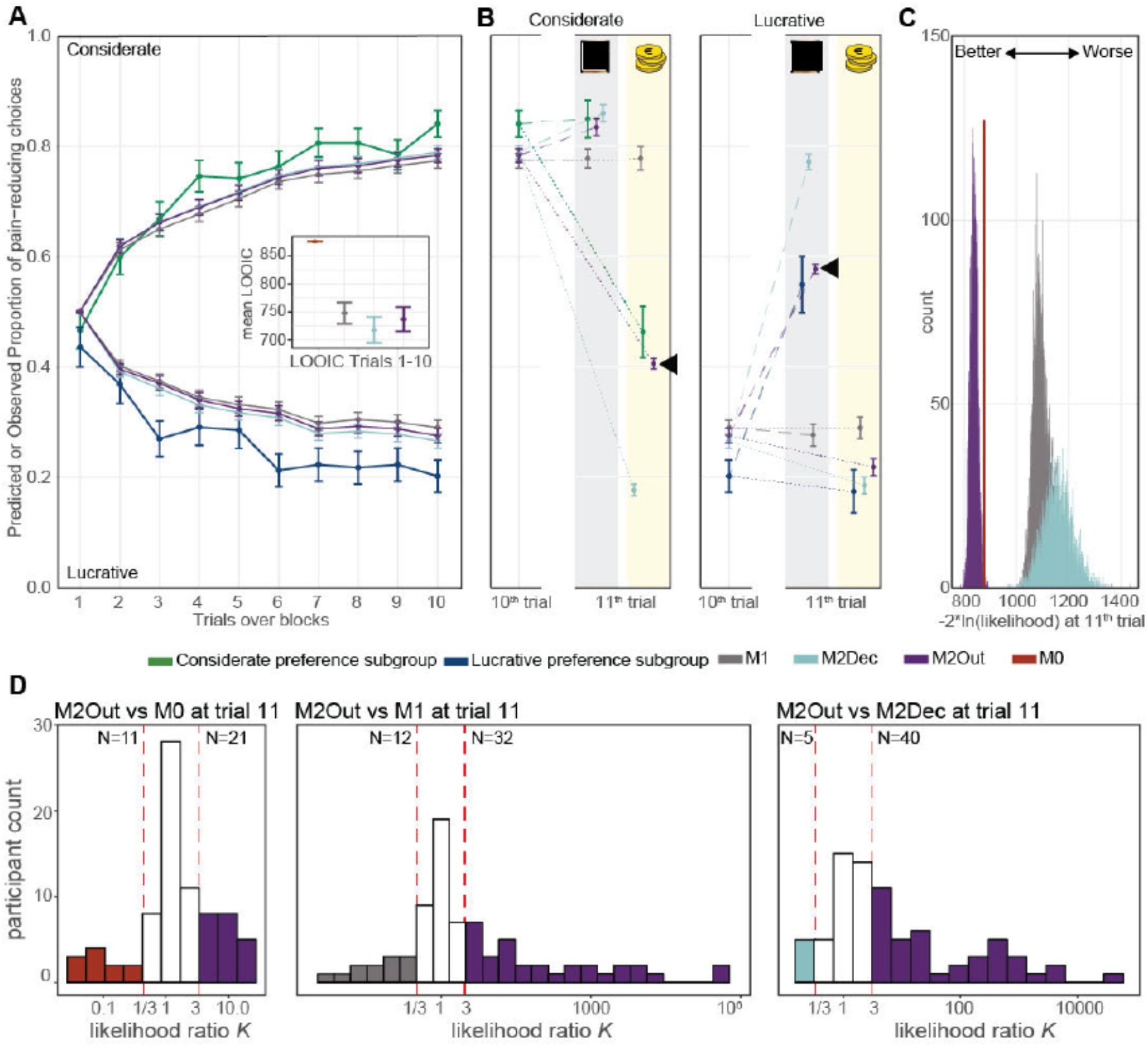
Model Comparison. **(A)** For the Conflict DropOut blocks of the online experiment, the graphs shows the trial-by-trial proportions of pain-reducing and lucrative choices for the first 10 trials of each block for participants with considerate (green) and lucrative (blue) preferences (preference classification is based on all conflict trials also including No-DropOut blocks), together with the predictions made by Ml. M2Dec and M2Out. MO always predicts 0.5 and is thus not shown, and participants with ambiguous preference are also not shown. In all cases, choices and predictions are averaged over all blocks (and samples for the models), and error bars then represent the s.e m. across participants. The models were fitted to the choices in the first 10 trials only. The inlet, shows the leave-one-out information criterion (LOOIC(33)) of the models over the first 10 trials, with the error bars representing the estimation error. MO performs poorly, but the three learning models perform similarly. **(B)** White background: participant choices and model predictions for the 10^th^ trial as in (a). Gray and yellow backgrounds: participant choices and model predictions for the 11^th^ trial, with dashed lines connecting the 10^th^ trial to 11^th^ trials in blocks in wliich money is removed (gray background), and the dotted lines those where shocks are removed (yellow background), separately for the considerate and lucrative preference groups. Note that the 11^th^ trial was not included in the fitting of the models. M2Out (marked by black arrowheads) makes the best predictions for the 11^th^ trial. **(C)** Likelihood of the choices during devaluation at the 11^th^ trial over all participants (incl. those with ambiguous preference) and including both money and shock Dropout trials, under all four models. For each of the 4000 posterior draws, we summed the log likelihood of the 11^th^ trial for all participants and blocks, and the histograms show the distribution of those likelihood over the 4000 posterior draws. Results were multiplied by −2 to place them on the information scale also used for LOOIC, that quantifies the information lost by each model, with lower values indicating a model with better predictions, and zero would indicate perfect predictions without lost information. M2Out clearly outperformed all other models in its ability to predict out-of-sample choices after devaluation of one of the outcomes. Note that we use LOOIC in (b), because these first 10 trials were included in the fitting of the model, but log-likelihood in (c) because it is not included in the model fit. **(D)** The distribution of likelihood ratio (*K*) in favour of M2Out across subjects relative to the competing models based on the log-likelihood of the 11th trial. *K*=exp(log_lik_M2Out)/exp(log_lik_OtherModel). *K>3* represents subjects with at least modest evidence to have been more likely to have used M2Out than the alternative, and *K*<⅓, more likely to have used the alternative model. Note that participants with *K*>3 always outnumber those for *K*<⅓, showing that also at the single subject level, M2Out is the model most favoured by the data during devaluation.

Figure 6C estimates the performance of the models at the group level, by adding the log-likelihood of all participants and trials, and plotting this overall likelihood for each of the 4000 samples drawn from the posterior distribution. To compare models at the single subject level, we extracted the mean log-likelihood of the 4000 samples for each participant and model, and calculated the likelihood ratio *K* in favour of M2Out, against each of the other models, as the ratio of their mean likelihoods. Using the traditional bounds for moderate evidence that have become standard in Bayesian statistics (*K*>3(34)), in all cases we find more participants with evidence in favour of M2Out than for the other models (Figure 6D).

That M2Out outperforms M2Dec in particular, is because M2Out scales expected values based on *wf*. For instance, an extremely considerate participant has *wf*≈0, and the EV_M_ thus remains close to zero for both symbols across the first 10 trials, because Out_M_ are multiplied with this *wf*≈0 to calculate PE_M_ (Figure 2), and EV_M_ thus is the sum of many PE_M_ close to zero. Accordingly, when shocks are removed, choices are between two symbols with EV_M_≈0, predicting the ~50% preferences observed. In contrast, for M2Dec, such scaling of EV by *wf* does not occur, and on the 11^th^ trial, participants have EV values for their less-favored outcome that are as differentiated as those of the favored outcome. Note that if we simulate a version of M2Dec that would maintain *wf* in the decision-phase of the 11^th^ trial, M2Dec would make predictions extremely similar to M2Out.

### M2Out can accurately recover the value of wf, and this value predicts both preference in our task and helping in a different task

For our winning model M2Out, fitting the model on the first 10 trials of the DropOut blocks leads to a wide range of *wf* values (Figure 7A) across participants that echoes the wide distribution of choices (Figure 5). Indeed the *wf* parameter estimates correlate highly with the proportion of pain-reducing choices (in both experiments, Kendall’s Tau_(wf,choice)_<-0.82, p<0.001, BF_10_>1000, Figure 7B). Parameter recovery further shows that if one simulates participants with different *wf*, M2Out accurately recovered much of that variance (r(*wf*_simulated_,*wf*_estimated_)=0.69, p<10^-6^, BF_10_>10^6^ Supplementary Figure 6). For our fMRI participants, we also performed a Helping task not involving learning that we had previously developed(4) (Supplementary material §3). In each Helping trial, participants viewed a victim receive a painful stimulation, and could decide to donate money. If no money was donated, the victim received a second stimulation of equal intensity. If some money was donated, each € donated reduced the second stimulation by one point on the 10 point pain scale. We found that the *wf* value estimated in our learning task in the fMRI experiment could significantly predict how much money participants on average donated to reduce the victims pain in this different Helping task (Kendall τ(wf,choice)=-0.47, BF_10_=76, p<0.001, Figure 7C), providing evidence that *wf* captures a property of the participant that generalises to other situations in which participants need to resolve a conflict between self-money and other-pain. A Bayesian multiple linear regression including *wf* together with more traditional trait questionnaires (four subscales of the IRI^26^ as a traditional measure of empathy and MAS^27^ as a traditional measure of the attitude towards money), found strong evidence that including *wf* improves the predictive power (BF_incl_=11.46, Supplementary Table 3), while none of these questionnaires explained variance in the Helping Task (all BF_incl_<⅓, Supplementary Table 3). This establishes the utility of our task as a predictor of costly helping. The distribution of the other parameters of M2Out can be found in Supplementary Materials §8.

**Figure 7.**
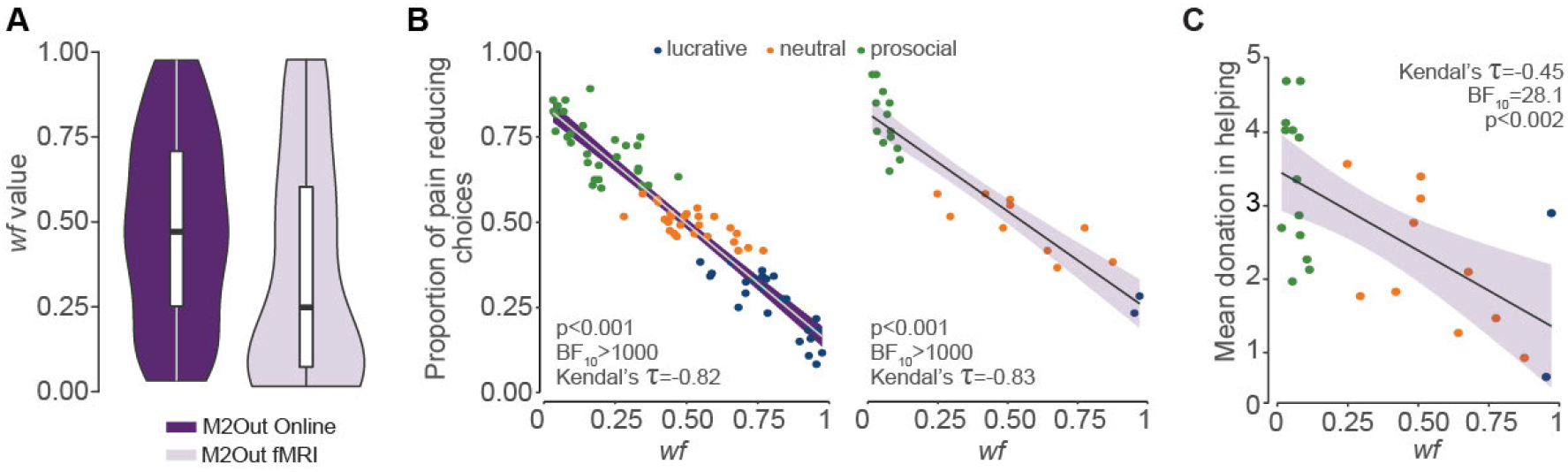
*Wf* Predictive power. **(A)** Distribution of *wf* parameter estimates for M2Out in the Online (purple, only including the first 10 trials of the Conflict-DropOut blocks) or fMRI (lavender, including all trials) across participants. The whisker plots represent the quartiles, dots outliers. **(B)** Relation between *wf* and proportion of pain reducing choices in the online (left) and fMRI (right) experiment. For the online experiment only the 10 first trials of DropOut blocks are considered in the model fit. The line and shading represent the regression line and its standard error, *p* and BF values are for a two-tailed test on the Kendall’s tau value. **(C)** Relation between *wf* in fMRI experiment and donation in the helping experiment.

### Neural correlates of prediction errors for shocks can be found in the pain observation network and the vmPFC

Participants showed robust brain activity across a wide network of brain regions when the outcomes of their decisions were revealed (Figure 8, Supplementary Table 5)

**Figure 8.**
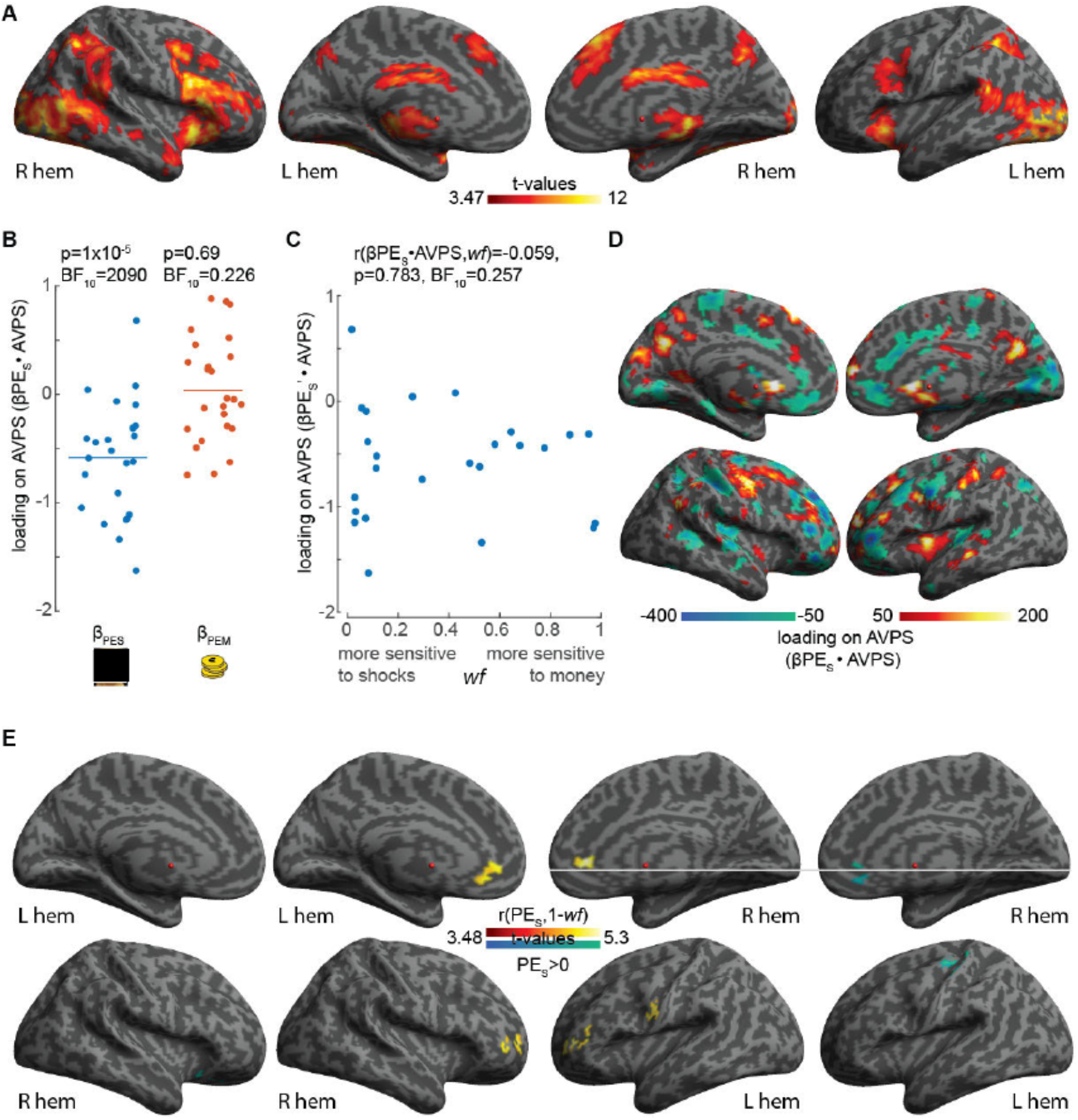
Key fMRI results. **(A)** Results of the contrast Outcomes>O indicating voxels where BOLD signals during the outcome phase (independently of prediction errors) are increased, p<0.001, k=FWEc=749 voxels. **(B)** Loading of the normalized parameter estimate images (β_PE_) for PE_S_ and PE_M_ on the AVPS(28). Each dot is one participant, the horizontal line is the mean. *p* and BF values reflect a one-sample parametric (because normality was not violated) *t*-test against zero in JASP. **(C)** Loading of the parameter estimate for PE_S_ (β_PES_) onto the AVPS as a function of *wf*. *p* and BF values represent the outcomes of a parametric correlation analysis using Pearson’s *r*. **(D)** Voxels contributing substantially to the loading of β_PES_ onto AVPS. Because the overall loading is calculated by first multiplying each voxel’s β_PES_ value with the value in this voxel of AVPS, and then summing these multiplication over all voxels, here we simply calculated the first step (the voxelwise multiplication of β_PES_ and AVPS), and averaged the result across our participants. To not overcrowd the image, we only show voxels with values above 50 or below −50. **(E)** Wann colors on the central two columns: regression between PE_S_ and 1-*wf*(p_unc_<0.001. k=FWEc=162voxels). This identifies voxels with signals that increase more for shock outcomes that are less intense than expected (i.e. positive PE_S_) in participants that place more weights on shocks (high 1-*wf* values). Cold color in the outer two columns: results of the contrast constant>0 in the same linear regression with (1-wf) (p_unc_<0.001, k=FWEc=132voxels). This identifies voxels with signals that increase with increasing PE_S_ after the variance explained by (1-*wf* is removed. All results in (E) were FWE cluster corrected at p<0.05, following cluster cutting at p_unc_<0.001. specified using the critical FWE cluster size FWEc. Note that the reverse contrasts for (e) did not yield results that survive FWE cluster correction. The light gray line helps the viewer to visually compare the location of the clusters across the two contrasts.

To examine whether signals in the affective vicarious pain signature (AVPS) covary with prediction errors for shocks (PE_S_), and whether they do so in a way that correlates with *wf* or not, we extracted the *wf*-normalized PE_S_ parameter estimate image (β_PES_) from each participant, and dot-multiplied with the AVPS neural signature. It is critical for the interpretation of our fMRI analysis that we use *wf*-normalized PE predictors in our fMRI analysis (as described in Methods: fMRI data analysis: PE *wf*-normalization) to avoid that *wf* influences results twice: in calculating the parameter estimate image and in calculating the regression of the parameter estimates and *wf*. The AVPS is known to have larger signals when viewing painful compared to non-painful facial expressions (28). We coded shock outcomes in terms of their *value* for a participant and not their intensity, i.e. a non-painful shock had value +1 and a painful shock of −1. Hence, trials with painful shocks should typically have negative prediction errors, and β_PES_ should load *negatively* on this neural signature. We found this to be the case (Figure 8B, t_(24)_=-5.55, p=1E-5, BF_10_=2090). Next, we asked whether the strength of the loading depended on *wf* using a Bayesian correlation, and found evidence of the absence of such a relationship (r=-0.058, p=0.78, BF_10_=0.257, Figure 8C). The facial pain observation network thus does carry signals that negatively covary with prediction errors for shocks in our paradigm, and these signals do not depend on personal preferences (*wf*). Voxels contributing substantially to the loading can be found in Figure 8D.

Next, we performed a more explorative voxel-wise linear regression that predicts the parameter estimate of the PE_S_ modulator using a constant and *wf*. As expected for a region involved in valuation, we found the vmPFC to have signals covarying positively with PE_S_ (i.e. higher signals when shocks were lower than expected). A more ventral cluster showed signals associated with PE_S_ in a way that depended on *wf* (Figure 8E, warm colors and Supplementary Table 6), while a more dorsal vmPFC cluster showed an association with PE_S_ after removing variance explained by *wf* (Figure 8E, cold colors and Supplementary Table 7). In addition, we find that the left somato-motor cortex, including BA4 and 3, also harboured signals covarying positively with PE_S_, with a more dorsal location representing it with a significant dependence on *wf*, and a more ventral region in a way that did not significantly depend on *wf*.

Because PE_S_ and PE_M_ predictors in our fMRI design are somewhat correlated due to the outcome structure of our task (average r=-0.26, ranging from −0.49 to −0.03 across our participants), we performed a parameter recovery analysis (See Supplementary Material §11) that confirmed that if we simulate participants with signals that covary with PE_S_ but not PE_M_, we recover significant parameter estimates for PE_S_ but not PE_M_, and vice versa. We also found that the PE_M_ parameter estimate images do not load on the AVPS, and found evidence of absence of such loading (t_(24)_=0.392, p=0.7, BF_10_=0.226, Figure 8B), confirming our ability to appropriately separate and localize PE_M_ from PE_S_ signals.

## Discussion

Our aim was to investigate how participants learn action-outcome associations during moral conflict - so called morally salient features (35) - that contrast gains for the self against pain for others. We had three central questions. First, whether such learning would be model-free or model-based, and whether they can be modeled using reinforcement learning models. Second, whether individual preferences for maximizing self-gains vs. minimizing other-pain would bias learning towards the preferred outcome. Third, if such bias exists, whether and how the brain would process other-pain exclusively in a biased fashion, or whether it might have co-existing representations that are biased and unbiased.

Our results indicate that participants’ choices appear to be dominated by biased model-based learning (16). The dominance of model-based learning was apparent (i) from the ability to provide above chance explicit reports for both outcomes that also captured the difference between Conflict and No-Conflict blocks and (ii) from the reaction to devaluation by not altering choices if the preferred outcome was preserved, but by switching preference when the preferred outcome was removed in the conflict blocks. That models were biased was borne out from the fact that (i) individuals had more differentiated reports for their favoured outcome-type (ii) that preferences after devaluation were asymmetric, with preference levels depending on the weight of the remaining quantity.

Bayesian model-comparison of our models confirmed this conclusion (see Supplementary Materials §12 for a detailed discussion of the predictions and rationale of each model). During the first 10 trials of Conflict, all the learning models outperformed the random-choice model (M0), but made virtually indistinguishable predictions. However, when one outcome was devalued, M2Out outperformed the other learning models at the group level, and in the majority of participants. Specifically, M2Out was the only model that captured the prevalent behavior of unchanged preference, when the less favoured outcome was removed and weak preference, when the most valued outcome was removed. This was because M2Out tracked separate self-money and other-shock expectations for each symbol (and is hence model-based), but did so in ways that depend on individual preference (and is thus what we call biased). For a minority of participants, we found evidence that unbiased model-based learning or model-free learning was a better description of choices under devaluation, and further studies may want to investigate which variables drive such individual differences.

That our participant’s learning seemed dominated by a model-based form of learning contrasts with a recent report that participants choices appear to be dominated by model-free learning when learning to prevent shocks to others(15). Two main experimental differences could account for these contrasting findings. Firstly, in Lockwood et al., agents following model-based learning do not outperform agents following a model-free strategy(15), while in our DropOut condition, model-based learning allows swifter switching to the most favourable option. This difference in incentives could have biased our participants towards the arguably more effortful model-based strategy. Second, in Lockwood et al., participants only had to track one outcome (shocks for self *or* others), while in our task, two outcomes (self-money and other-shocks) were always involved. Dual, conflicting outcomes may prioritise more model-based forms of learning. Overall, this difference emphasizes the need and opportunity to investigate how situational demands shape the choice participants make between model-based and model-free learning in morally relevant situations.

Considering the dominance of the biased model-based learning, we next asked how this biased model-based learning would be implemented in the brain. Given the considerable literature on learning to choose options that maximize gains for the self (22, 36), we focused on how shocks to others are processed in the brain under conflict. In principle, there might be two ways in which the bias in our model-based learning could be implemented in the brain: (1) the bias occurs so early that a person maximizing self-gain fails to process shocks altogether (but see Supplementary material §15 for preliminary eye-tracking data) or (2) all participants initially process shocks in similar ways and modulate these signals with *wf* only in later stages of processing. In case (1) we would fail to find BOLD signals associated with PE_S_ in a way that does not depend on *wf* anywhere in the brain, while in case(2) we could expect that earlier stages of processing (e.g. the network associated with pain observation(24–26, 28)) has signals associated with PE_S_ that do not depend on *wf*, while later stages of processing associated with valuation, particularly in the vmPFC(14, 22, 29–31) might have PE_S_ signals that depend on *wf* and inform choices. The psychological (37) and neuroimaging(38–42) literature cannot adjudicate these alternatives, as they have shown biases of empathy at both early and late levels of processing. However, our fMRI data speaks to option 2: the pain observation network, as quantified using the affective vicarious pain signature(28), had signals that covaried with prediction errors for shocks with evidence for the absence of a linear dependence on *wf*, while within the medial prefrontal cortex, we found more dorsal regions with signals that covary with prediction errors for shocks with magnitudes that did not significantly depend on *wf*, as well as more ventral signals that did. That the ventral but not the dorsal cluster had valuation signals that depend on *wf*, i.e. on whether a participant focuses on minimizing other-pain or not, dovetails with a finding of Nicolle et al. (32) that if participants have to switch from trial to trail between focusing on making a monetary decision according to their own values or those of someone else, more ventral mPFC voxels had valuation signals corresponding to the values of the current focus, while more dorsal clusters had valuation signals corresponding to those of the alternative focus.

A limitation of our fMRI study is that, unlike prediction errors for shocks and money that we show to be distinguishable, outcomes and prediction errors were so highly correlated (shock: mean r= 0.84, ranging 0.75-0.87; money: mean r= 0.84, ranging 0.78-0.86) that we cannot establish whether signals are truly representing prediction errors rather than outcomes. While this problem is not unusual (e.g. (12)), a novel version of the paradigm better dissociating prediction error and outcome will be necessary to explore whether the pain observation network simply processes outcomes or already integrates outcomes with expectations as predictive coding theories might predict(43–45). However, whether the pain observation network processes outcomes or prediction errors, it does so in ways that do not depend on *wf*, thereby providing evidence against the notion that the bias is so pervasive that the brain does not at all represent shocks independently of bias. This hybrid representation of some signals that are biased and others that are not, echoes the fact that in our Online experiments, even participants with lucrative preference have above chance access to the association of the symbols to shocks (Figure 4B), while the magnitude in probability difference does correlate with *wf* (Figure 4D). A promising direction in which our study could be expanded would involve the use of electro-physiological recordings. In particular, one could then explore whether signals in the pain observation network covarying with prediction errors earlier than the vmPFC signals that do depend on *wf*, and whether coherence across these regions increases during shock observation, to test the idea that vmPFC valuation may be informed by unbiased activity in the pain observation network.

In addition to these core fMRI findings, we made a couple of additional, less expected findings in our explorative analysis. First, we failed to find signals robustly associated with prediction errors for money when applying a whole brain correction for multiple comparisons (but for a lower uncorrected threshold results see https://osf.io/rk8w4/?view_only=98b193a58aff48dda40b9d3d91ac5254 Figure8_fMRI/SecondLevel_Regression_PEM_wf). That we had few participants with very high *wf* (i.e. with highly lucrative preferences) and more participants with low *wf*, in our fMRI study, could help explain why it was more difficult to find signals that represent the self-money compared to the other-shocks. Additionally, money was always delivered (i.e. money was delivered also in the low-money condition), and we had robust activations in regions associated with monetary rewards when outcomes were revealed. The strength of that trial-independent signal may have made it more difficult to detect the apparently smaller differences between trials with higher and lower monetary outcomes. In the future, it might be helpful to either compare conditions with no monetary reward at all against a condition with monetary reward, and to recruit a larger number of lucrative participants. Second, we also found regions around the left central sulcus encoding prediction errors for shocks in ways that depend on *wf* or not, extending into both SI (area 3b and 1) and MI (area 4). While it is tempting to interpret these activations as representing some form of embodiment of the somatic pain of others onto the observer’s own somatomotor cortex (4, 27), such conclusions should be made with care considering the limitations of reverse inference, in particular given the explorative nature of this whole brain analysis (46). Neuromodulating these foci, akin to what we have done previously in our helping task (4), and exploring the impact on choices, would be important to explore their causal contribution to learning and decision making.

Participants varied substantially in their preferences under conflict. In our Online study, about one third prefered the symbol leading to high monetary gain, another third, the symbol minimizing the pain to the other, while for the remaining third the number of trials we acquired did not suffice to identify a significant preference. This pattern was true despite that to maximize conflict, in the online experiment we asked each participant to adjust the amount of money on offer for the high money outcome to be equivalent to the cost of the other receiving the painful shock-intensity we used (Supplementary Material §2). Indeed, participants that had a lower indifference point, later choose the lucrative option more often, and these individual differences in choice are thus not a trivial consequence of individual differences in the amount of money on offer. In our models, we find that an individual parameter, the weighting factor (*wf*, conceptually similar to the salience factor alpha in the Rescorla-Wagner Learning Model (19)) is an effective way to capture these individual differences in the relative weight placed on the money vs shocks. Importantly, *wf* had external validity, in that it predicted how much money the same participants gave to reduce shocks to the same confederate in a different task, the Helping Task (4), which does not require learning. We also found that *wf* was better than our trait measures of empathy and money attitude scales at predicting donation in this Helping Task. Recent work has supported the view that state empathy is regulated by motives and context (see (37, 47) for reviews). Our moral conflict task creates financial incentives known to downregulate empathy (37). It is thus perhaps unsurprising that the IRI, which measures self-reported trait empathy and which does not probe empathy under such conflictual situations, fails to predict decisions under conflict, while *wf*, which is estimated for learning during moral conflict, is better at predicting decisions during moral conflict in the Helping task. This speaks to the need to develop situated state empathy measures that probe the propensity of participants to deploy their empathy in specific situations, to complement measures of empathy as a context free trait (37, 47). Our paradigm might be particularly well suited in phenotyping antisocial populations and could provide insights into the atypical tuning of their morally relevant learning and decision-making in the spirit of computational psychiatry (48). This is because psychopathic offenders show atypical instrumental learning (49) and reduced activity in empathy related networks while witnessing the pain of others (50) despite typical levels of trait empathy in self-report questionnaires.

In our fMRI study, most participants (13/27) showed a significant preference for the pain-reducing option, with only 3/27 showing a significant preference for the lucrative option. Why this distribution was different from the Online experiment is unclear, but two factors may play a role. First, in the fMRI experiment, that was conducted on site, participants were under the impression that all shocks were delivered in real time to the other person, while in the online experiment, they were led to believe that a subset of the shocks would later be delivered to another person. This may have made the value of reducing shocks higher in the fMRI than in the Online experiment. Second, the vicinity of the experimenter in the fMRI experiment may have increased the motivation to act in a socially desirable way compared to the online experiment. As there is no significant correlation between our variables of interest (*wf* and percentage of considerate choices) with age, the difference in age between the experiment is unlikely to have caused this difference.

Our study has several additional limitations. First, we limited our model comparison to a number of hypotheses driven by RLT models. We did not test ratio or logarithmic ratio models of valuational representation in this study. These valuational structures are known to occur but are less often indicated in modeling gains and losses to the self (17). Future experiments could be optimized to explore whether such alternative ways to combine these values may be more appropriate under certain moral conflicts. Second, our participants or at least some of them, may not use a RLT model at all. Instead, they may use rules such as ‘choose a symbol randomly, and only switch if you encounter *x* unfavorable outcomes in a row’. Future studies may benefit from studying how well such heuristics may perform. Third, we used *wf* to address quantitative individual differences. We recognize that *wf* only indicates preferences during our tasks and that it is risky to interpret the *wf* as suggesting stable moral values, as is the case for all behavioral studies absent strong evidence regarding stable moral commitments. Future studies may wish to explore whether different individuals may be best captured using qualitatively different models and in conjunction with validated evidence of long-term moral commitments and values. This might be particularly relevant when including participants with independently demonstrated morally considerate commitments on the one hand, or psychiatric disorders affecting social functioning on the other. Fourth, our evidence for a biased model-based approach hinges on drop-out trials in our Online experiment, and our request for probability reports. We introduced these trials to rigorously reveal the nature of the learning that participants deploy in conflict situations - and they indeed provided the data necessary to adjudicate across our models. However, we must also consider the possibility that these very trials also influenced participants to deploy the model-based learning that provides more optimal decisions during drop-out and more accurate probability reports. Performing a conflict task with only a single devaluation trial on a large number of participants may be a way to exclude this possibility. However, if participants were to have adapted their strategy to optimally fit the requirements of the task, M2Dec would have been even more adaptive.

To summarize, our data sheds light onto the processes at play when adults have to learn that certain actions lead to favorable outcomes for us but harm others, while alternative actions are less favorable for us but avoid harm to others. We show that in our task, the choices of a majority of participants are best described by a biased model-based approach, in which some brain signals (e.g. in the pain observation network) covary with prediction errors for the harm to others in ways that do not depend on whether participants prefer to maximize gains for self or minimize harm to others, while others (e.g. in the vmPFC) do depend on this individual preference. We foresee that the neurocognitive model we introduce and this task will be particularly useful, in the future, to understand how neurocomputational processes may differ in antisocial populations that often fail to make choices that appropriately consider harm to others in situations in which such harm results from pursuing personal goals.

## Materials and Methods

Two independent experiments were performed: an Online study and an fMRI study. Behavior was tested in a larger sample of participants, but this had to be done online, using the Online platform Gorilla (https://gorilla.sc/), due to COVID-19 restrictions in place at the time. Brain activity was measured in a smaller number of participants in our fMRI scanner. Table 1 gives an overview of the number of participants and experimental conditions included in each study.

### Participants

In total, 106 healthy volunteers with normal or corrected-to-normal vision, and no history of neurological, psychiatric, or other medical problems, or any contraindication to fMRI (for the fMRI experiment only) were recruited for our experiments (Table 1). Particularly, 27 participants (37y±17SD; 27f) took part in the fMRI, and 79 (25y±7SD, 39f) in the Online experiment, with the two samples being independent. Participants taking part in the Online experiment were significantly younger than the ones who took part in the fMRI study (Mann-Whitney test, BF_10_=8.022, p<0.001). However, choices (in terms of percentage of considerate choices) and *wf* did not significantly depend on age (Kendall’s tau(age, %Considerate choices)=0.113, BF_10_=0.543, p=0.096; Kendall’s Tau(age,wf)=-0.115, BF_10_=0.561, p=0.091). Participants for the online task were recruited through Prolific (https://prolific.ac/), while those for the fMRI through advertisements of the experiment on social media.

As two of the 27 participants in the fMRI were left handed, and stimuli showed movements of the right hand of the actor, to reduce variability possibly induced by lateralization of the brain responses, these two participants only performed the tasks off-line (i.e. no fMRI data acquired). The studies were approved by the Ethics Committee of the University of Amsterdam, The Netherlands (2017-EXT-8201, 2018-EXT-8864, 2020-EXT-12450). Consent forms for participation and authorization for the publication of images have been obtained.

### Online Experiment general procedure

In the Online experiment, participants’ main task was the probabilistic reinforcement learning tasks, in which participants had to choose between two new symbols based on learning how much money each symbol could earn them, and whether the symbols lead to a painful shock to another participant (Figure 1). The task included a deceptive cover-story in which participants were informed, at the very beginning of the study, that while they perform the task online, a second participant is present in our lab. It was explained that the separation of the two participants allowed the experiment to be performed under COVID19 restrictions. In reality, no one was invited to our lab, and this cover story served to create a situation in which participants took what they thought to be decisions with real implications for self and others, while at the same time minimizing discomfort to others (no shocks actually delivered to the other person). Participants then received an general overview of all the tasks they will encounter during the experiment. The text also informed them that 15 trials would be randomly selected from the main Learning task, and that the self-money of these trials would be added to the online participants payment, while the electrical stimulation associated with these trials would then be delivered on the hand of the other participant in the lab. Participants knew they would be offered the choice to see (via Zoom), or not, the other person receiving the stimulation at the end of the task. The experimental session started with a Stress Tolerance questionnaire assessing participants’ susceptibility to stress, which generated the advice to continue or not with the experiment (Supplementary material §13). No participant received the advice to withdraw, and the session continued with the Optimization Task, aimed at determining the amount of money necessary to create a true conflict for a given person between self-gain and other-shock (this value is referred to as *indifference point*). This was done by asking participants to choose between different predetermined combinations of money and stimulation intensity (Supplementary material §2). The choices in this task also count toward the final bonus and number of stimulation to the other. The experiment then continued with the core Reinforcement Learning Task (see below). The experiment ended with a debriefing about the deception, asking how much they believed the cover story (Supplementary material §4) and with a series of questions to investigate participants’ motivations and their feelings toward the experiment (Supplementary material §14). Participants were aware that they could withdraw from the experiment at any time.

### Reinforcement Learning task (Figure 1-D)

#### Conflict

Participants had to learn symbol-outcome associations under a moral conflict - i.e. in a context in which choices that are better for the self (high monetary gain) are usually worse for the other (painful shock), while those that are worse for the self (low monetary gain) are usually better for the other (non-painful shock). Each block was composed of 10 trials (Figure 1A) in which participants had to learn the outcomes associated with each of the two presented symbols. The same symbol pair was presented in all trials of a block, but changed across blocks. In each block one of the symbols would most likely lead to higher monetary reward for the participant, and a noxious electrical stimulation to the dorsum of the actor’s hands (lucrative symbol or choice), and the other symbol to lower monetary reward and a non-noxious stimulation to the actor (pain-reducing symbol or choice). Symbol-outcome associations were drawn independently for each symbol and outcome, following the schema illustrated in Figure 1B. This procedure helped decorrelate representations of shock and money. While the monetary reward was presented in the outcome screen as a numerical amount of euros, the outcome for the other was presented as a video recording of the author A.N. while receiving the stimulation (Supplementary material §1). Participants could therefore infer whether the stimulation was or not noxious only based on the facial reaction of the actor in the videos. Participants were aware that the person in the video and the other participant in the lab were not the same individual, and that the videos were pre-recorded to illustrate what responses to the stimulations would look like.

#### No-Conflict

This condition was identical to the Conflict blocks with the exception that the symbol-outcome associations did not likely introduced a moral conflict because the symbol that was usually best for the self was also usually best for the other: one symbol was most likely associated with high monetary reward for the participants and non-noxious stimulation to the other participant, while the other symbol was usually associated with low monetary reward and a noxious stimulations. There was therefore a clear incentive to choose for the symbol leading to the higher reward, as it was also the symbol that would be most beneficial for the other.

#### ConflictMoneyDropout

Each block included two sets of 10 trials. The first 10 trials were identical to those presented in the Conflict condition. After the 10th trial, the participants were informed that the money outcome was removed and the block continued with the last 10 trials, in which participants would only see the videos of the actor receiving the stimulation, and no money would be rewarded. Participants were informed that the probability of each symbol of being associated with high or low stimulation would remain the same over the 20 trials.

#### ConflictShockDropout

This condition was identical to the ConflictMoneyDropout with the exception that the removed outcome was the stimulation to the other. From the 11th to 20th trials participants therefore only saw the amount of money associated with the chosen symbol and no consequences were to happen to the other.

#### No-ConflictMoneyDropout and No-ConflictShockDropout

These two conditions were the same as the ConflictMoneyDropout and ConflictShockDropout, except that the first 10 trials were the same as those presented in the NoConflict conditions, in which symbol-outcome associations did not likely introduce a moral conflict.

The position of each symbol on the screen within a pair was randomized across trials. Symbols composing a pair were kept the same across individuals (i.e. the same two symbols formed the same pair throughout the experiment), but pairs were randomly distributed across conditions and participants. Symbol-outcome associations were determined based on the matrix in Figure 1B, and were kept constant across participants. Conditions were presented in a randomized order within and across participants.

#### Explicit recall questions

At the end of each of the four Conflict and four NoCoflict blocks only, participants were asked, for each symbol separately, to recall the probability of that symbol to be associated with higher monetary reward and with the noxious stimulation using a scale from 0 to 100% (Figure 1C). The order in which the two symbols were presented and the type of question (association with high shock or with high reward) was randomized between participants. The slider starting position was always at 50%.

### fMRI Experiment

Participants in the fMRI experiment performed two experimental tasks: a Helping task as published in Gallo et al. (4), and the Conflict condition of the reinforcement learning task used in the Online experiment. The experiment consisted of four sessions of the Helping task and one session of the Conflict task, positioned after the first two sessions of Helping. fMRI results of the Helping task will be the object of a separate publication. In the current work we only report the fMRI data associated with the Learning task, and the amount of money participants were willing to give up to reduce the intensity of the confederate’s stimulation in the Helping task in order to test the predictive power of the weighting factor.

Briefly, the Helping task showed videos of another participant (author S.G.) receiving noxious stimulations of different intensities (Supplementary material §3). Participants were then endowed with 6 euros, which they could decide to keep for themselves or to give all or in part up to proportionally reduce the intensity of the next stimulation. Videos either showed the facial reaction of the confederate or the hand reaction (no face). Because the online experiment mainly showed facial reactions, only donations associated with the trials showing the facial reactions in the Helping task were included in the current manuscript (results would though not change when including the full donation dataset (Kendall τ(wf,choice)=-0.54, BF_10_=13.19, p=0.004). Participants were led to believe that the videos they saw were a life-video-feed of another participant receiving these shocks in a closeby room, although in reality they were prerecorded movies. All participants were presented the same videos, albeit in randomized order, so that we can compare the average donation of the participants as a willingness to give up money to reduce the pain of others.

As the participants of the fMRI experiment came in person to the lab, the cover story was slightly different from that adopted in the online version. For the fMRI we used the same cover story used and validated in Gallo et al. (4). Each participant was paired with what they believed to be another participant like them, although in reality it was a confederate, author S.G. They drew lots to decide who plays the role of the learner (or the donor in the Helping task) and of the pain-taker. The lots were rigged so that the confederate would always be the pain-taker. The participant was then taken to the scanning room while the confederate was brought to an adjacent room, connected through a video camera. Participants were misled to think that electrical stimulations were delivered to the confederate in real-time, and that what the participants saw on the monitor was a live feed from the pain-taker’s room. In reality, we presented pre-recorded videos of the confederate’s reactions.

In contrast to the Online experiment, in which the high-money amount was selected for each participant using the Optimization task, in the fMRI Learning task, the high-money amount offered was fixed at 1.5€ for all participants, and corresponded to the average amount associated with the indifferent point in the Optimization task (1.53 ± 0.37SD; Supplementary Material 2). The low-money amount was the same as in the Online experiment, 0.5€.

At the end of the fMRI tasks, participants were debriefed and asked to fill out the interpersonal reactivity index (IRI) empathy questionnaire (51), and the money attitude scale (MAS) (52). To assess whether participants believed that the other participant really was receiving electrical shocks, at the end of the experiment, participants were asked ‘Do you think the experimental setup was realistic enough to believe it’ on a scale from 1 (strongly disagree) to 7 (strongly agree). All participants reported that they at least somewhat agreed with the statement (i.e. 5 or higher).

The fMRI task was programmed in Presentation (www.neurobs.com), and presented under Windows 10 on a 32inch BOLD screen from Cambridge Research Systems visible to participants through a mirror (distance eye to mirror: ~10cm; from mirror to the screen: ~148cm). The timing of the task was adapted to the requirements of fMRI: each trial started with a jittered fixation cross lasting 3-9 seconds. Then the two symbols appeared and participants could make their choice without a time restriction. After the button press, there was a fixation cross raging from 3-9 seconds and the video with a duration of 2 seconds followed.

### Analysis of Behavioral Data

Quantification of choices was performed in RStudio (Version 1.1.453) and Matlab (https://nl.mathworks.com/). Statistical analyses were then performed using JASP (https://jasp-stats.org, version 0.11.1), to provide both Bayes factors and *p* values. Bayes factors were important because in many cases we are as interested in evidence of absence than in evidence of the presence of an effect, and p-values cannot quantify evidence for the absence of an effect (53). For instance, we expect that removing money outcomes in considerate participants should not alter choices, while removing shocks should. We used traditional bounds of BF_10_>3 to infer the presence of an effect and BF_10_<⅓ to infer the absence of an effect (34, 53). Two-tailed tests are indicated by BF_10_,i.e. *p*(Data|H_1_)/*p*(Data|H_0_) while one-tailed tests are indicated by BF_+0_. Where ANOVAs were used, we report BF_incl_ which reports the probability of the data given a model including the factor divided by the average probability of the data given the models not including that factor. Normality was tested using Shapiro-Wilk’s. If this test rejected the null-hypothesis of a normal (or bi-variate normal for correlations) distribution, we used non-parametric tests such as the Wilcoxon signed rank or Kendall’s Tau test, while if the null-hypothesis was not rejected, we used parametric tests. We always used default priors for Bayesian statistics as used in JASP.

### Computational Modeling of Behavioral Data

Our experiment represents a variation of a classical two armed-bandit task and was modeled using a reinforcement learning (RL) algorithm with a Rescorla-Wagner updating rule (19). We compared 4 models explained in Figure 1E. Models were fitted in RStan (version 2.18.2, http://mc-stan.org/rstan/) using a hierarchical Bayesian approach, i.e. by estimating the actual posterior distribution through Bayes rule. Our models were adapted from the R package *hBayesDM* (for “hierarchical Bayesian modeling of Decision-Making tasks”) described in detail in Ahn et al (2017) (54).

Model comparison was also performed in a fully Bayesian way. All the 3 models were fit just on the conflict condition, and just the first 10 trials of each block (the ones involving the conflictual choices). To assess the ability of a model to fit data of these initial 10 trials that were included in the parameter-fitting, we use the leave-one-out informative criterion (LOOIC (33)), which computes a pointwise log-likelihood of the posterior distribution to calculate the model evidence, rather than using only point estimates as with the other methods, e.g. of Akaike information criterion (AIC (55)) and the deviance information criterion (DIC (56)). The LOOIC is on an information criterion scale: lower values indicate better out-of-sample prediction accuracy. For the Conflict-Dropout blocks, on the other hand, we aimed to assess the ability of a model fitted on the first 10 trials of a Conflict-Dropout block to predict behavior on the devaluation trial (trial 11). The model was then fitted using the first 10 trials of all the Conflict-Dropout blocks, and no leave-one-out procedure was necessary to assess the predictive performance on the 11th trial because it was not included in the parameter fitting. We thus directly used the log likelihoods of the 11th trial, which capture the probability of the choices on the 11th trial given the model fitted on the preceding 10 trials. Specifically, at the group level, we added the log likelihood of all the 11th trials of all the participants for each of the 4000 draws from the posterior distribution, and then examined the distribution of these values (Fig. 4C). To compare the ability of different models to predict this 11th trial at the single subject level, we then calculated a likelihood ratio (similar to a Bayes factor) for each participant: we summed the log-likelihood of the 8 11th trials available for each participant, and used the mean across the 4000 draws as our point estimate. We then calculated the ratio of the exponents of the log-likelihoods for the competing models.

As priors on the hyperparameters □ and *wf*, we use the recommended Stan method of using a hidden variable distributed along N(0,1), that is then transformed using the cumulative normal distribution to map onto a space from 0 to 1. For □, we use the same method but then multiply the result by 5 to have the function map onto the interval [0,5].

For the Dropout blocks of the Online experiment, in which one of the two outcomes was removed after the 10th trials, the three models M1, M2Out, M2Dec were modified from the 11th trial to account for the fact that participants were told which quantity would be removed. For the M2 models, this was implemented by setting EV=0 for the removed quantity before decision-making on the 11th trial. In addition, *wf* is modified only to value the remaining EV (i.e. *wf*=1 if shocks are removed and *wf*=0 if money is removed). For M1, we cannot reset expectations for a specific quantity (shock or money) and EV has to remain unchanged. However, the *wf* is adapted (i.e. *wf*=1 if shocks are removed and *wf*=0 if money is removed), which maximizes what can be learned from the following trials. In all cases, the outcomes for the missing quantity are always set to zero after Dropout.

### MRI Data acquisition

MRI images were acquired with a 3-Tesla Philips Ingenia CX system using a 32-channel head coil. One T1-weighted structural image (matrix = 240×222; 170 slices; voxel size = 1×1×1mm) was collected per participant together with an average of 775.83 EPI volumes ± 23.11 SD (matrix M x P: 80 x 78; 32 transversal slices acquired in ascending order; TR = 1.7 seconds; TE = 27.6ms; flip angle: 72.90°; voxel size = 3×3×3mm, including a .349mm slice gap).

### fMRI Data preprocessing

MRI data were processed in SPM12 (57). EPI images were slice-time corrected to the middle slice and realigned to the mean EPI. High quality T1 images were coregistered to the mean EPI image and segmented. The normalization parameters computed during the segmentation were used to normalize the gray matter segment (1mmx1mmx1mm) and the EPIs (2mmx2mmx2mm) to the MNI templates. Finally, EPIs images were smoothed with a 6mm kernel.

### fMRI Data analysis

For the Learning Task, our aim was to identify brain activity that scales with PE_S_ for shock when the outcome is revealed, and to compare this activity across participants with different weighting factors. Analyses and the experimental design therefore focused on the outcome phase. In line with other studies (e.g., (12)), activity during the decision phase will not be analysed here for two reasons. First, to isolate activity when outcomes are revealed, we randomized the interval between outcomes and the response screen. As a result, decisions can have occurred at any time between the last outcome and the next button press, making it difficult to capture the activity linked to that decision. Second, the button press required for the response triggers significant brain activity in frontal regions that could have been hard to dissociate from the valuation processes we are interested in.

To analyse activity during the outcome phase, we ideally would have included predictors for the monetary outcome (Out_M_) and the shock (Out_S_) as well as their prediction errors (PE_M_ and PE_S_). Unfortunately, PE_S_ and Out_S_ are too highly correlated (0.741, ranging 0.750 to 0.866 across our 27 participants), and the same is true for PE_M_ and Out_M_ (0.749, ranging 0.781 to 0.862 across our 27 participants), to allow us to include them into a single design matrix. Indeed, we performed a parameter recovery attempt that simulates voxels with various mixtures of Out_S_, PES, Out_M_ and PEM across 24 participants with the addition of noise, and tried to recover them using design matrices with all 4 parametric modulators, and failed to have appropriate recovery (data not shown). This is not surprising, because for time-efficiency, here we aimed to focus on the initial learning of the association, including only 10 trials per block, and during the initial learning outcomes with high Out_S_ will also cause high PE_S_ and trials with high Out_M_ will also cause high PE_M_. Our parameter recovery simulations however confirmed that we can include PE_S_ and PE_M_ in a single design matrix, as these are only weakly correlated (average r=-0.26, ranging from −0.49 to −0.03 across the participants), and correctly identify voxels simulated to contain PE_S_ but not PE_M_, PE_M_ and not PE_S_ or both (Supplementary Materials §11). The parameter recovery also indicated that we can dissociate voxels in which participants with lower *wf* show stronger PE_S_ related signals. This then allows us to focus on the processing of shocks in the context of learning by identifying voxels with signals that covary with PE_S_ in ways that do or do not linearly depend on *wf*, while being aware that signals that covary with PE_S_ might in fact have covaried with Out_S_.

Our fMRI design matrix thus included the following regressors. The decision regressor started with the appearance of the two symbols and ended with the button press of the participant. The outcome regressor was aligned with the presentation of the video and had a fixed duration of 2 seconds, corresponding to the duration of the stimulus. The decision regressor had 5 parametric modulators (EV_S1_, EV_S2_, EV_M1_, EV_M2_, choice) and the outcome regressor 2 parametric modulators (PE_S_ and PE_M_). The modulators were derived from the winning M2Out, with EV_S1_, EV_S2_, EV_M1_, EV_M2_ representing the expected values in terms of shock and money for symbol 1 and symbol 2, and choice had value 1 or 2 based on the chosen symbol. Symbols ‘1’ or ‘2’ correspond to the lucrative and pain-reducing choice respectively.

#### PE wf-normalization

Because we are interested in whether PE_S_ or PE_M_ representations depend on *wf* or not, we divided PE_S_ with (1-*wf*) and PE_M_ with *wf* before entering them into the design matrix. As a result, the first PE_S_ value in the parametric modulator would always be PE_S_=-1 if it was a high-shock outcome or PE_S_=+1 for a low-shock outcome, independently of the participant’s *wf* value. If signals covary with PE_S_ in a way that does not linearly depend on *wf*, the parameter estimate across participants (β_PES_) would violate H_0_:β_PES_=0, but not H_0_:r(β_PES_=0,wf)=0. If signals covary with PE_S_ in a way that does linearly depend on *wf*, it would violate both of these H_0_. Note that for outcomes, the coding was +1 for good outcomes (i.e. high money or low shock) and −1 for bad outcomes (i.e. low money or high shock). EV and PE follow that polarity.

Results were then analysed in two ways. First, to improve reverse inference, we use a multivariate signature developed to distinguish the sight of facial expressions of pain vs. neutral facial expressions that captures the overall activation of the facial pain observation network as a single scalar value: the affective vicarious pain signature (AVPS(28)). To explore if signals in this network covary with PE_S_, we then simply dot-multiplied the PES parameter estimate volume for each participant separately with the AVPS, after having brought the AVPS into our fMRI analysis space using ImageCalc. The result of the dot-multiplication indicates how much the covariance with PE_S_ loads on the AVPS. We then brought these values into JASP, and compared them against zero and correlated them with *wf*. Because the values were normally distributed, we used parametric analyses. We did the same for PE_M_ to confirm that our analysis can separate PE_M_ and PE_S_ signals. Second, we performed a similar analysis at the voxel level, by bringing the parameter estimate images for PE_S_ into a second level linear regression using a constant and wf as the two predictors. A t-test on the constant then reveals regions in which signals covary with PE_S_ after removing variance explained by *wf*. A t-test on the *wf* parameter estimate then reveals regions where the signals covary with PE_S_ in ways that depend on *wf*. To test if signals covary with 1-*wf*, we simply used a negative contrast in the t-test. Results were familywise corrected at the cluster level using the established two-step procedure in SPM: (i) for cluster-cutting we visualized results at *p*_unc_ < .001 k=10, and identified the FWEc minimum cluster size for family wise error correction from the results table, (ii) we reloaded the results at *p*_unc_ < .001 k=FWEc, so that all displayed results survive FWE at cluster-level.

## Data Availability

All data and analysis code are available at https://osf.io/rk8w4/?view_only=98b193a58aff48dda40b9d3d91ac5254

## Acknowledgements

The work was funded by European Union’s Horizon 2020 research and innovation programme under ERC-StG ‘HelpUS’ 758703 to VG, Dutch Research Council (NWO) VIDI grant (452-14-015) to V.G and VICI grant (453-15-009) to C.K. M.S. thanks the John Templeton Foundation (Grant 21338) for support in developing the ideas regarding self/other valuational representation. We thank C. Gavanozi, I. Gembutaite and B. Hoekzema for helping with data acquisition in the Replication and Outcome-Dropout study. We thank A. Veggerby Lind for helping recording the stimuli used in the Replication and Outcome-Dropout studies. We thank A. Gentile for his input and help with developing the learning models. VG, CK, AN, LD, KI, SG, LF and RP thank P. Lockwood as her work inspired the development of the learning tasks.

## Author contribution

The experiments were conceived by VG and CK with input from all authors. Funding and project leadership by VG. The fMRI data were collected by KI with the help of SG; the behavioral data by LF. Pilot data for the preparation of the study were obtained by RP, AN and SG. FMRI data were analyzed by KI with guidance from V G and CK; the RLT models were developed and programmed by AN, CK, LD, LF, MS, NE, and VG. AN, CK, KI, LF, MS, and VG wrote the manuscript with edits and comments from all other authors.

## Supplementary material

### 1. Stimuli creation and validation for the Learning task

#### Videos

Videos were generated in house. The authors SG and AN played the role of the actress during the fMRI and Online experiment respectively. All videos were recorded to start with an initial neutral facial expression, followed by the facial expression in response to the electrical stimulation delivered to the right-hand dorsum. The upper part of the actress’ body was clearly visible on a black background. The actresses were encouraged to produce realistic and clear facial expressions in response to the temporally unpredictable stimulations. The intensity of the stimulation was decided with the actresses, and the facial expressions were exaggerated to reflect a clear expression of pain. All original recordings were cut to last 2s and to have the frame showing the beginning of the facial expression in response to the stimulation exactly at 1s. All videos were validated by independent groups of subjects, who were asked to rate the intensity of the pain experienced by the confederate on a scale from 1 to 10, with ‘1’ being ‘just a simple touch sensation’ and ‘10’ being ‘most intense imaginable pain’. As low pain intensity videos we selected the ones with ratings of 1 and 2 and as high pain intensity videos we selected the ones with ratings of 4, 5 and 6 Videos were edited using Adobe Premiere Pro CS6 (Adobe, San Jose, CA, USA).

The videos used for the Learning task in the fMRI experiment were selected from the Hand videos recorded for Gallo et al., 2018 (1) (https://elifesciences.org/articles/32740, see also Supplementary Material§3 below, and https://doi.org/10.7554/eLife.32740.006, https://doi.org/10.7554/eLife.32740.009). Thirteen videos of value 1 and 14 of value 2 were used as low pain stimuli, while 9 videos of value 4, 13 of value 5 and 3 of value 6 were chosen for as high intensity stimuli. A full description of how these videos were created can be found in (1). A similar procedure was then used to generate the video of the Online experiment (for two examples see https://osf.io/rk8w4/?view_only=98b193a58aff48dda40b9d3d91ac5254_Figure1_Task). The main difference was that in the stimuli for the Online experiment both actress’s hands were shown (Figure 1A), instead of only the right hand. For the Online experiment, an initial pool of 300 videos showing a painful stimulation and 300 videos showing an innocuous stimulation were recorded. The final pool of stimuli was then selected based on an online validation (survey done through Gorilla, https://gorilla.sc/) performed by 191 participants (aged 18 – 35, 103 females). Participants were recruited through Prolific (https://prolific.ac/). From the initial pool, 140 high intensity and 130 low intensity stimuli were selected. Since for each condition we needed 140 trials (120 trials with both outcomes plus 40 trials with money dropout), 10 low videos were repeated twice in each condition. The exact same videos used in the Conflict blocks were repeated in the No-Conflict blocks, this allowed us to rule-out any effect due to the difference in the displayed stimuli between conditions. The repetition did not have any negative impact on participants’ performance, since in the Online experiment participants were aware of the fact that these videos had been pre-recorded.

The high intensity pain stimuli used in the fMRI Learning task had an average rating of 4.76 ± 0.66SD and the low intensity stimuli of 1.52 ± 0.51SD (Bayesian Mann-Whitney t-tests W=675, p<0.001,BF_10_=2710.5). For the Online study, the high intensity pain stimuli had an average rating of 5.74 ± 0.47SD and the low intensity stimuli of 1.54 ± 0.18SD (Bayesian Mann-Whitney t-tests, W=18200, BF_10_=7.7*10^11^, p<0.001). There was no difference between the average rating of the low intensity pain stimuli used in the fMRI and in the Online experiment (Mann-Whitney t-test W=1750, BF_10_=0.24, p=0.983). On the other hand, the difference between the average rating of the high intensity pain stimuli used in the two experiments was significant (W=502.50, BF_10_=456.31, p<0.001). Having high intensity pain stimuli perceived as of lower intensity in the fMRI experiment might a priori predict more lucrative preferences in the fMRI compared to the Online experiment. In contrast, lucrative preferences were rarer in the fMRI than in the Online experiment (Table 1), suggesting that this difference in the high intensity stimuli did not impact behaviour substantially.

#### Symbols

All the symbols used in the learning task were created using Adobe Illustrator. For the creation of symbols we used simple geometrical shapes without obvious meaning. Each symbol was then paired with a second one created using the same shape elements organized in a different position. All the symbols covered the same area on the screen. Twenty-four pairs of symbols were used in the test: 12 pairs were used in the Conflict, and 12 in the No-Conflict condition. The absence of *a priori* preference for a symbols in each pair was assessed in an independent group of subjects (Bayesian Wilcoxon Signed-Rank test, W=31, BF_10_=0.31, p=0.547).

### 2. Optimization task

In the Online experiment, before the learning tasks, we determined the amount of monetary reward that would have a subjective value equivalent to the painful shock received by the other participant. This task enabled us to personalize the amount of money participants were later offered as high reward in the learning tasks to create a meaningful conflict. Participants always had to choose between a pain-reducing option combining 0.5€ for them with a low shock to the confederate, and a lucrative option combining a higher amount of money for themselves with a high shock to the confederate. The amount of money offered in the lucrative option varied in steps of 0.25€ across the 5 types of choices (see the table below). Before making the choice, the participant was able to see the intensity of the electrical stimulation for the confederate by playing pre-recorded videos.

**Supplementary Table 1.** Optimization Task. The table lists the five possible options offered during the optimization task, with the number of participants to which each option was assigned during the Learning Task of the Online experiment. The numbers in brackets indicate the number of participants showing a Lucrative (L), Considerate (C) or Ambiguous (A) preference during the task.

|  | Pain-reducing option | Lucrative option | Nr participants |
| --- | --- | --- | --- |
| 1 | 0.5€ + Low Shock | 1€ + High Shock | 12 (6L, 3A, 3C) |
| 2 | 0.5€ + Low Shock | 1.25€ + High Shock | 22 (12L, 8A, 2C) |
| 3 | 0.5€ + Low Shock | 1.5€ + High Shock | 13 (3L, 7A, 3C) |
| 4 | 0.5€ + Low Shock | 1.75€ + High Shock | 7 (2L, 2A, 3C) |
| 5 | 0.5€ + Low Shock | 2€ + High Shock | 25 (1L, 6A, 18C) |

Each of the 5 types of choices were presented 6 times, for a total of 30 decisions. A sigmoid was then fitted to the choice data, and the indifference point was selected based on where the sigmoid crosses the 0.5 pain-reducing proportion. To select the high reward to offer during the learning task, we picked the lowest value for which each participant chose half of the times the low-electrical stimulation option and half the high electrical stimulation option. In case there was not a value for which the number of pain-reducing choices was equal to the number of lucrative choices, we picked the lowest value for which the number of lucrative choices was higher than the number of pain-reducing choices. If a participant had always chosen the pain-reducing option, we picked the highest value (2€) as the high reward for the learning experiment. Conversely, if a participant had always chosen the lucrative option, we picked the lowest value (1€) as the high reward for the learning experiment. Due to an error in programming the task, to 4 participants who had 1€ as real indifference, 1.25€ was offered as high reward during learning. Another subject whom should have been offered 1€ got offered 2€ as high reward. This could have induced a more lucrative behaviour in this subject, however this participant was still in the Considerate group, meaning that this didn’t have a strong impact on performance. In the Online experiment, the average indifference point was 1.53 ± 0.37SD, which matched the 1.5 € given as high reward to every participant in the fMRI experiment.

### 3. Helping Task

In the fMRI experiment, participants performed the Helping Task presented in Gallo et al. (1) as main experimental task. Only the behavioral results (i.e. average donation per participant associated with the Face stimuli) of the Helping Task are included in the current publication. Briefly, participants performed 60 trials in which they watched a first (prerecorded) video of the same confederate as in the Learning Task receiving a painful stimulation. The intensity of the stimulation could vary between 1 and 6 on a 10 point pain scale, and was chosen on each trial by the computer program. In each trial participants also receive 6 credits, and could decide to donate some of these credits to reduce the intensity of the second stimulation to the confederate. Each credit donated back to the experimenter reduced the next stimulation by 1 point on the 10 point pain scale. Participants then watched a second video showing the confederate’s response to the second stimulation. At the end of the task, participants were paid the sum of the amount of money that they had kept for themselves from all the trials divided by 10. We capture individual differences as the average number of credits given up per trials (“donation”). Two types of videos were presented. One showing the confederate receiving an electroshock on the hand and expressing the pain she felt by only reacting with facial expression. The other showed a belt hitting the dorsum of the confederate right hand, and the confederate expressing how much pain she felt by a reaction of the hand alone. The face was not visible in the latter stimuli. Hand and Face videos were presented in separate sessions.

In total there were 2 sessions of 15 trials each presenting the hand video and 2 presenting the face. The single session with 6 blocks of Learning task was presented at the end, after the four Helping Task sessions. This was because the fMRI experiment was centered on the Helping task, and the Learning task was meant as a pilot data, which should have been followed by a second fMRI data acquisition centered to the Learning, which was however impossible due to COVID19 restrictions.

**Supplementary Figure 1.**
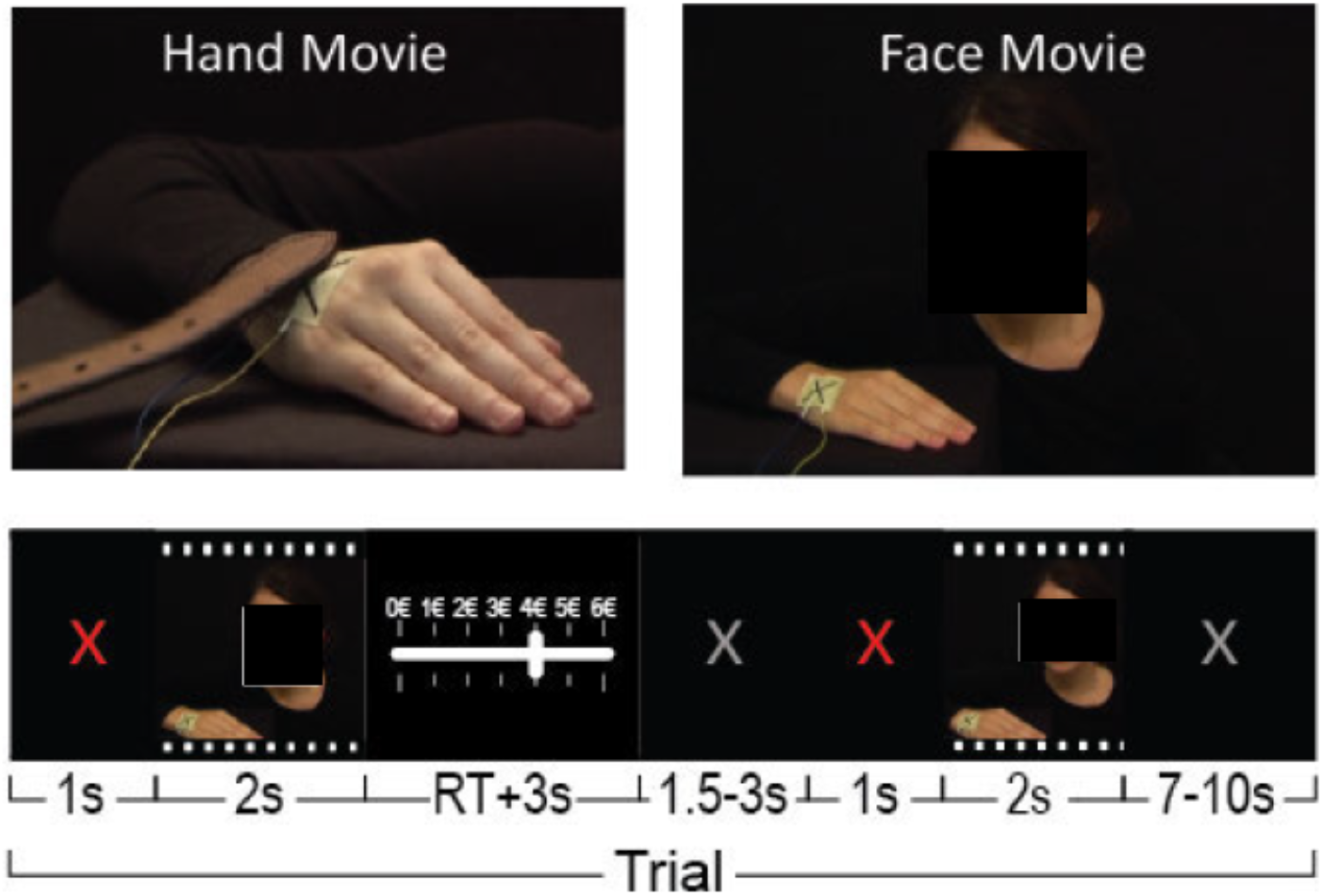
Helping task structure. Top: two screen shots taken from one exemplar video showing the confederate receiving the slap on her right hand, and from a video showing the confederate receiving the shock on the hand and manifesting its painfulness through facial reactions. Bottom: trial structure.

### 4. Participants’ belief that someone will get shocks

At the end of the Online study, participants were asked to express the degree of agreement to the statement: “Before choosing whether or not to see the other person receiving the electrical stimulations, I believed that someone was really going to receive them” on a scale from 1 (strongly disagree) to 7 (strongly agree). If they at least somewhat disagreed, they were additionally asked whether they nevertheless acted as if they believed someone would get shocks or not. An ANOVA with level of agreement as a nominal factor showed that average choices were not influenced by how much a participant believed in shock delivery (Main effect of belief in shocks, F_(6,72)_=0.739, p=0.62, BF_incl_=O. 109). However, of the 28 Non-Believers (i.e. those who responded ‘somewhat disagree’, ‘disagree’, or ‘strongly disagree’), 24 reported they nevertheless acted as if someone were to receive the shocks (Supplementary Table 2). This might explain why people that doubted anyone would get shocks would nevertheless give up money to reduce shocks. In contrast, the 4 participants reporting they acted as if no one were to get shocks (red in Supplementary Figure 2) chose pain-reducing options on average significantly less than the other 75 participants (Independent sample test t_(77)_=-2.97, p=0.004. BF_10_=8.75). In the fMRI study, all 27 participants reported that they believed that someone was getting a shock.

**Supplementary Table 2.** Distribution of participants’ beliefs. The number in each cell is the participants count. Believers were considered those reporting values from 4 to 7 (neither agree to totally agree), non-believers those reporting 3 to 1 (somewhat disagree to totally disagree). Nr=number.

|  | Total Nr of participants | Believers | Non-Believers |  |
| --- | --- | --- | --- | --- |
|  |  |  | *Acting as if believing* | *Non-acting as if believing* |
| Online participants | 79 | 51 | 24 | 4 |
| fMRI participants | 27 | 27 | 0 | 0 |

**Supplementary Figure 2:**
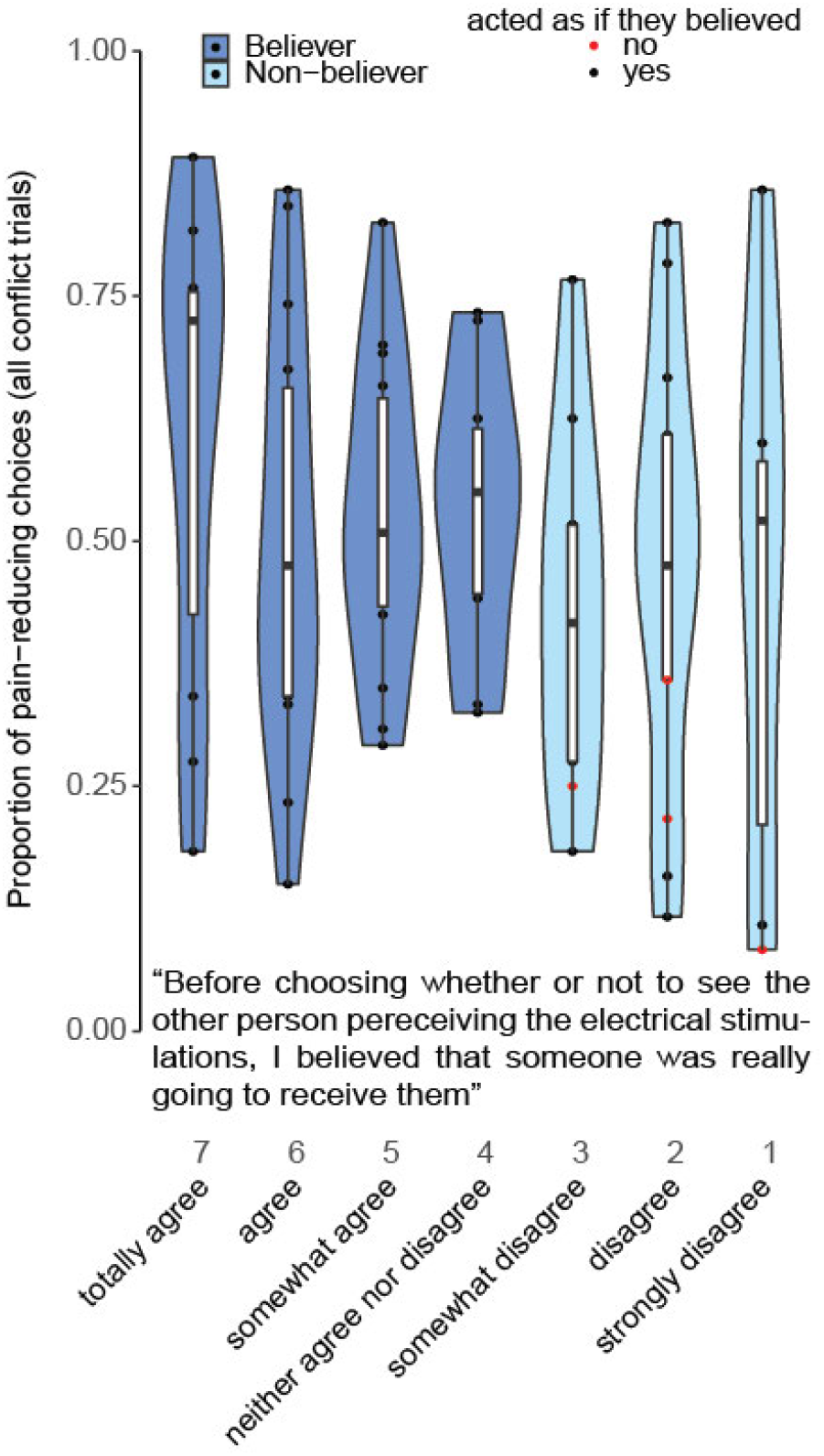
Effect of belief in the fact that the victim will receive shocks on choice allocation in the Online study. The figure shows the proportion of pain-reducing choices across all the conflict trials of the Online experiment (including No-Dropout and the 10 first trial of Dropout blocks) as a function of the answer provided. Whisker plots represent the quartiles, black dots the outliers, and red dots participants reporting that they did not act as if someone were getting shocks. Participants responding that they at least somewhat disagree are considered Non-Believers for the purpose of Supplementary Figure 5.

### 5. Participants’ motivation

To better understand what drove participants’ choices in the Online learning task, at the end of the Online task, participants were asked the question “What motivated your choices?”. Five options were proposed to the participant and they could choose to select 1 or more. In the case they selected more than 1, they were asked to select them in order of importance, starting from the most important. The options presented to participants were the following:

A. Gaining as much money as possible
B. Giving a good impression
C. Being in line with my moral beliefs
D. Avoiding to see the other person receiving the shocks
E. Preventing harm to others

Supplementary Figure 3A below reports how often a particular motivation was chosen as the primary motivation in each group.

**Supplementary Figure 3.**
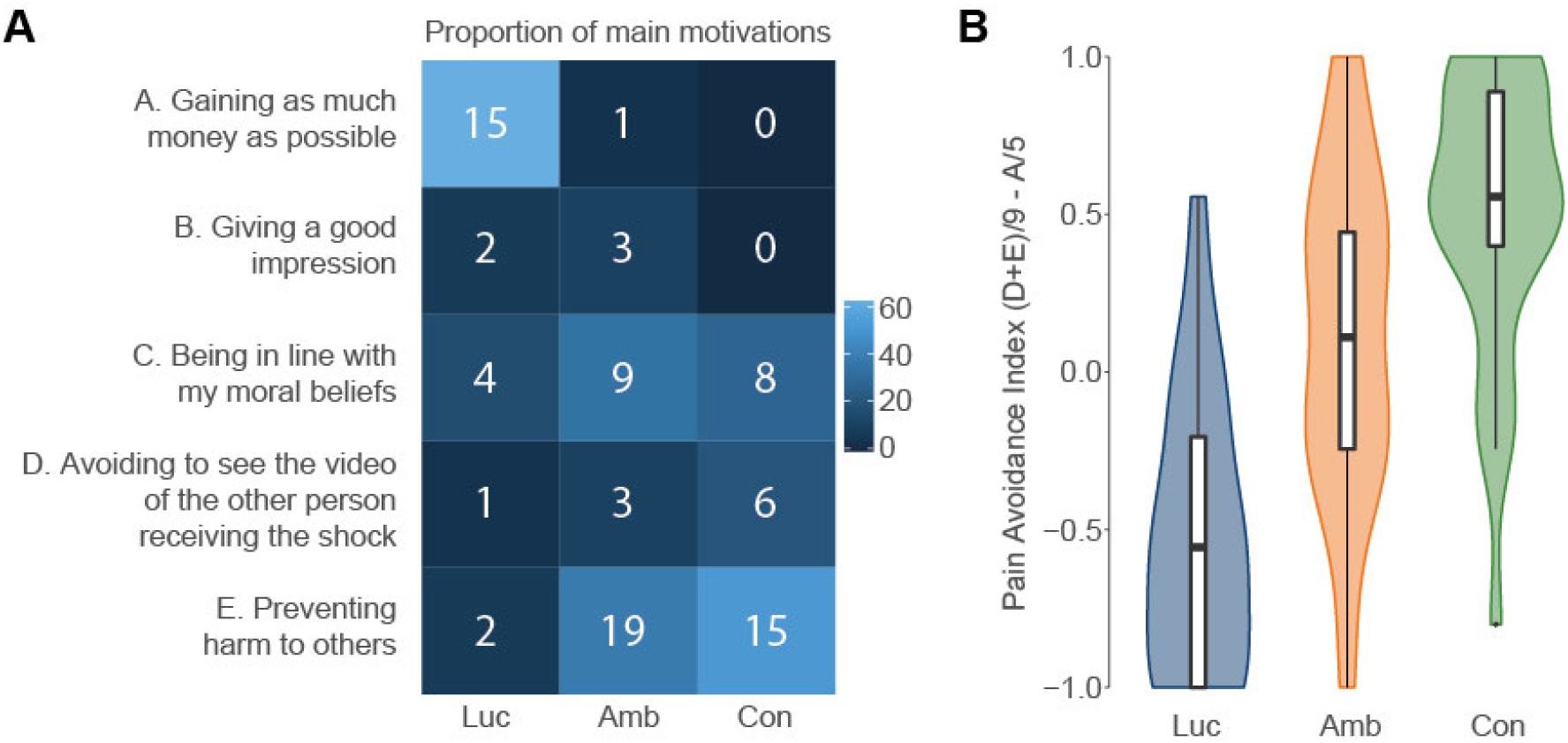
Motivation as a function of preference. **(a)** for each of the preference groups, the color indicates the proportion of participants that have chosen a particular motivation as their primary motivation, and the white numbers represent the actual number of participants in that cell. **(b)** The value of the pain-avoidance index, calculated as (D+E)/9-A/5, as a function of group preference. The whiskers represent the quartile. This index has the value of +1 if participants select pain avoiding motivations as their first two motivations (i.e. preventing harm to others and avoiding to see the video of the other person receiving the shocks) and do not select ‘gaining as much money as possible’, and −1 if they select ‘gaining as much money as possible’ and not the two pain avoiding motivations.

Since there were 5 alternatives, to quantify motivation, values from 0 to 5 were assigned to A-E according to its position in the list of motivations (0 if it had not been selected, 5 if it had been indicated as first motivation, 4 as second, 3 as third, 2 as forth and 1 as fifth). To assess whether the motivation overall was more lucrative or pain-reducing, we then derived a secondary measure of ‘*Pain-Avoidance*’ that would be negative if the motivation was mainly ‘gaining as much money as possible’ and positive if the motivation was ‘preventing harm to others’ or ‘avoiding to see the other person receiving the shocks’. Specifically, this was calculated as (D+E)/9-A/5. We divided D+E by 9 and A by 5, because a person with maximum pain reducing motivation could choose D and E as their first motivations and thus obtain 5+4=9, and one maximally lucrative would choose A as their first motivation. We decided on this contrast of D+E and A, because we found D+E to correlate highly negatively with A (tau=- 0.38, p<0.001,BF_10_=29232.57). Supplementary Table 3 below gives an overview of the average pain avoidance score calculator for the Lucrative, Considerate and Ambiguous group preferences separately (see also Supplementary Figure 3B).

**Supplementary Table 3:** Descriptive statistics for the Pain-Avoidance score. As can be seen, the score indeed varies all the way from −1 to 1 amongst our participants, but is not normally distributed, and is thus analysed non-parametrically.

| | Pain-Avoidance score $((D+E)/9-A/5)$ | | |
| --- | --- | --- | --- |
|  | lucrative | ambiguous | considerate |
| Participants | 24 | 26 | 29 |
| Mean | -0.519 | 0.120 | 0.506 |
| Std. Deviation | 0.447 | 0.509 | 0.440 |
| Shapiro-Wilk | 0.903 | 0.960 | 0.887 |
| P-value of Shapiro-Wilk | 0.025 | 0.384 | 0.005 |
| Minimum | -1.000 | -1.000 | -0.800 |
| Maximum | 0.556 | 1.000 | 1.000 |

To test whether participants that chose the pain-reducing options during the task, within or below chance (i.e. considerate, neutral and lucrative) differed in their pain avoidance score, we performed a Bayesian ANOVA with group as factor. This analysis revealed a significant main effect of group (Kruskal Wallis tests, H_(2)_=35.18, p<0.001,) and post-hoc Bayesian Mann-Whitney t-tests confirmed differences between all the 3 groups (Considerate-Ambiguous: W=196.50, p=0.002, BF_10_=9.31; Lucrative-Ambiguous: W=111.50, p<0.001, BF_10_=38.14; Lucrative-Considerate: W=47.50, p<0.001, BF_10_=397.01).

Testing each value against zero using a Wilcoxon signed rank test showed that for the considerate preference group values were significantly above zero (W=380, p<0.001, BF_10_=714.32), for the lucrative preference group they were below zero (W=13.5, p<0.001, BF_10_=748.88) but for the ambiguous preference group, they were close to zero (W=227.5, p=0.19, BF_10_=0.48). This suggests that participants in the Considerate and Lucrative group could and decided to consciously report motivations that were in line with their choices, while the Ambiguous group appeared to have less clearly polarized monetary or pain-avoiding motivations.

### 6. Choices in the NoConflict Dropout and across all 20 trials

**Supplementary Fig. 4.**
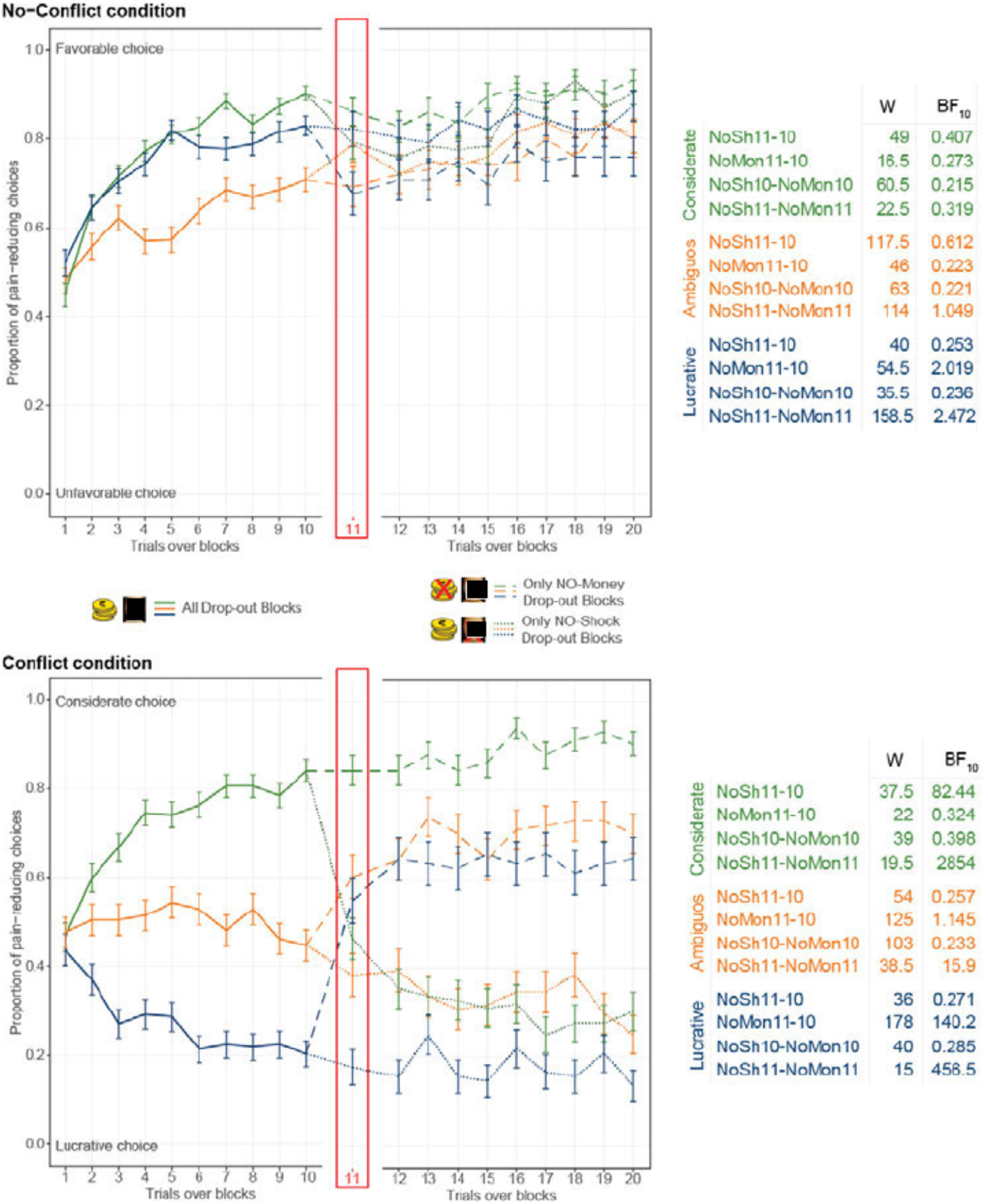
Participant’s choices in the DropOut blocks. For both Conflict and No-Conflict blocks, the figure show’s the average ± sem pain-reducing choices as a function of trial separately for the three preference groups. Note that preference grouping is based on all Conflict trials, including the No-Dropout blocks, while the figure only show’s choices on the Dropout trials. Tables summarize the Wilcoxon signed-rank values and BF_10_ for key comparisons. BF rather than p values are shown, to specify evidence for (BF_10_>3) or against (BF_10_<⅓) the presence of a difference. Here we also show’ choices on trials 12-20 after Drop-out. These choices are less relevant to the main paper, because they capture what is probably a different form of learning, w’hen symbols are only associated with 1 outcome rather than 2 outcomes that can conflict. NoSh=No-Shock, i.e. the outcome shock has been removed, and only money remains. NoMon=No-Money. i.e. the outcome Money has been removed, and only the shock remains.

### 7. Model comparison during Conflict Dropout excluding certain participants

**Supplementary Figure 5 (Related to Fig 4).**
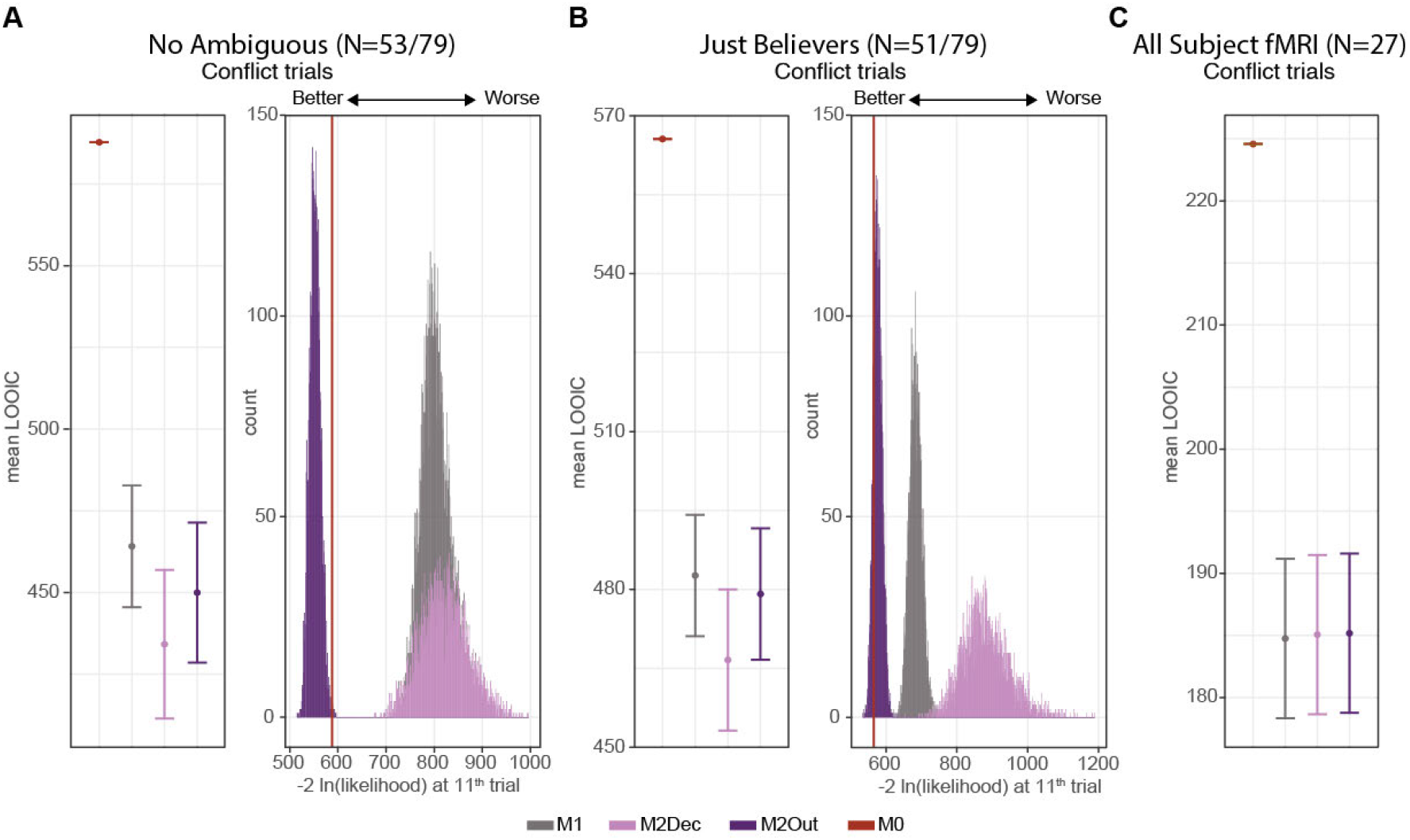
Model Comparison. (A) Mean LOOIC and the Likelihood of the choices during devaluation at the 11^th^ trial (as inlet Figure 6A, and Figure 6C), but excluding participants with ambiguous preferences. (B) As in (A) but excluding participants that expressed doubts. (C) Mean LOOIC for all fMRI participants.

### 8. M2Out Parameter Distributions and Recovery

The distribution of the parameters that were estimated for ***τ, λ***S and ***τ, λ***M can be found in Supplementary Figure 6A,B. We also performed a parameter recovery assessment on the M2Out model to assess the robustness of inferences on the parameter values estimated from this model. Generating values for the parameters *wf*, ***λ***S and ***λ***M were all drawn from a uniform distribution between 0 and 1, and ***τ*** from an exponential distribution between 0.2 and 5 (i.e., the natural logarithm of ***τ*** was drawn from a uniform distribution between −1.61 and 1.61). Parameter sets were sampled using a Latin hypercube sampling design (2, 3) to ensure an adequate coverage of the joint parameter space. The simulated data were generated using the full process of the M2Out model (i.e., simulated trial 1 informs simulated trial 2, etc.), with the amount of data matching the fMRI experiment (i.e., 6 blocks, each with 10 trials). Parameter values were estimated using Differential Evolution Markov Chain Monte Carlo (4), with 12 chains run in parallel for 2000 iterations and the first 1000 iterations discarded as burn-in. The priors on the *wf*, ***λ***S and ***λ***M parameters were all truncated normal distribution (between 0 and 1) with mean 0.5 and standard deviation 0.2, while the prior for the ***τ*** parameter was a truncated normal distribution (between 0 and infinity) with mean 1 and standard deviation 3. The estimates plotted in Figure 6c display the estimated posterior means from each simulated data set.

In the main manuscript, we use a hierarchical Bayesian model to estimate these parameters under the assumption that we sampled multiple individuals from the same underlying population. In particular for ***λ***S and ***λ***M, it is likely that different participants may gravitate onto similar ***λ***S and ***λ***M given the similarity in volatility experienced by them. For the parameter recovery we perform here, this assumption is not true: we deliberately sampled the entire space of possible ***λ***S and ***λ***M using a uniform distribution. Accordingly, for the parameter recovery, each simulated behavior was analysed individually using a non-hierarchical model. The priors for this individual implementation were not informed by the hyper-parameters of the final study, and were simply aimed at informing us about the proportion of all possible variance in these parameters that can be retrieved from analysing participants one at a time. The inverse temperature (***τ***) parameter had a relatively broad distribution across our participants in our two experiments (Supplementary Figure 6A), and parameter recovery shows that when simulating participants with different ***τ***, the ***τ*** estimates recovered by fitting M2Out correlated quite highly with the simulated ***τ*** values (Kendall’s ***τ*** (***τ***_simulated_, ***τ***_estimate_) 0.53, p<0.001, BF_10_>1000). As reported in the main text, the same was true for *wf* In contrast, parameter recovery shows that although the learning rates can be significantly recovered by M2Out, the correlation values are more modest than for *wf* and ***τ*** (Kendall’s ***τ*** (***λ***S_simulated_, ***λ***S_estimate_)=0.25, Kendall’s ***τ*** (***λ***M_simulated_, ***λ***M_estimate_)=0.24, both p<0.001, BF_10_>1000), and the ***τ*** values obtained from our model-fitting should thus be interpreted more tentatively. In particular, we find that ***τ*** is difficult to estimate for the outcome that participants consider less in their choices: for simulated participants with *wf*<0.1 (i.e. that minimize shocks to others), Kendall’s ***τ*** between simulated and estimated values drops to 0.03 for ***λ***M but is 0.36 for ***λ***S, while for simulated participants with *wf*>0.9 (i.e. that maximize gains to the self), it is 0.32 for ***λ***M but drops to 0.06 for ***λ***S. Accordingly, we did not include these parameter estimates in the main manuscript due to their reduced robustness.

**Supplementary Figure 6.**
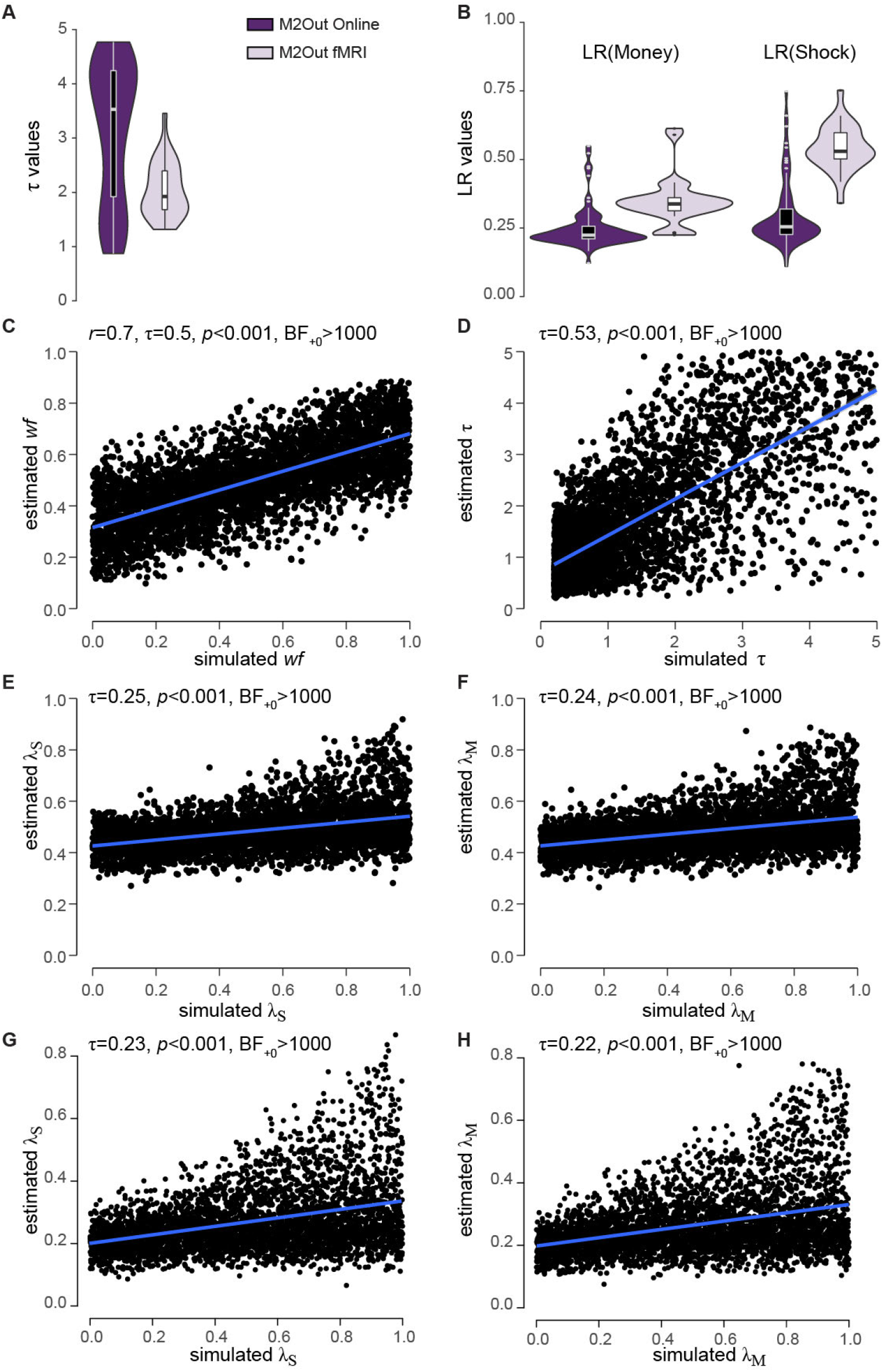
**(A, B)** Distribution of parameter estimates for M2Out in the Online (purple, only including the first 10 trials of the Conflict-DropOut blocks) or fMRI (lavender, including all trials) across participants. The whisker plots represent the quartiles, dots outliers. **(C-F)** Parameter Recovery Performance of a non-hierarchical implementation of M2Out. The figure shows the result of simulating the choices of 4000 participants that use M2Out for their decision, with a range of *wf*, ***τ***, ***λ***S and ***λ***M values, then estimating the parameters using M2Out. See §8 for details of the procedure. Note that this was done non-hierarchically to avoid influence from one simulation on the other, using priors that were not informed by the findings of our actual study. We performed a Shapiro-Wilk test for bivariate normality that considered for each of the four parameters both the simulated and estimated values. For *wf*, p=0.048, and we thus report both Pearson’s r and Kendall’s τ. For the other three parameters, the Shapiro-Wilk p was always below 0.001, and we thus only report Kendall’s τ. The *p* and BF values always refer to one-tailed tests with H1: ***τ*** >0, as this test does not require bi-variate normality. Note how for *wf* and ***τ***, the estimated values capture the simulated values fairly well, although the slope of the regression line is <1. For ***τ***, however, the prior distribution (truncated normal distribution, between 0 and 1, with mean 0.5 and standard deviation 0.2) appears to have a stronger influence on the estimated values than the simulated data, which strongly compresses the estimated values towards the peak of the prior and reduces the coupling between simulated and estimated values. **(G-H)** Adapting the priors for ***λ***S and ***τ***M to 0.2, closer to the values observed in the Online study, shifts the recovered parameters to a range closer to what was observed in the online study.

It might also be noted, that the median learning rates estimated by the hierarchical Bayesian model for the Online experiment (Supplementary Figure 6B) was close to 0.25, and was very different from the medial of the parameters recovered by the non-hierarchical model with a prior with mean 0.5. Adjusting the prior of the parameter recovery to match the hyperparameter obtained in the Online experiment fixes this anomaly.

### 9. Predicting helping based on *wf* but not IRI or MAS

**Supplementary Table 4.** Bayesian linear regression posterior summary of coefficients. The table summarizes the Bayesian linear regression model comparison for models explaining the donation in the Helping task using the *wf* of the Learning task, the subscales of the IRI (5) (FS, PT, EC and PD) and MAS (6) (Power-Prestige, Retention-time, Distrust and Anxiety). The first column indicates the variable under consideration, followed by the mean and sd of the regression parameter estimate, followed by the prior probability of including each variable (p(incl)) and the posterior probability of including each variable given the data (p(incl|data)). BF_incl_ indicates how-much more likely models including a variable are compared to the average of those not including this variable. BF_incl_>3 is considered moderate, and BF_incl_>10 strong, evidence that a variable explains donation. BF_incl_<⅓ indicates moderate evidence against a variable explaining donation. While shows a BF_incl_>5, all other variables have a BF_incl_<0.7. The most likely model given the data is therefore the one including the *wf* alone (7).

|  | Coefficient | Mean | SD | P(incl) | P(incl\|data) | BF inclusion |
| --- | --- | --- | --- | --- | --- | --- |
|  | Intercept | 2.643 | 0.211 | 1.000 | 1.000 | 1.000 |
| | $wf_{10T\_M2Dec}$ | -1.767 | 1.075 | 0.500 | 0.835 | **5.058** |
| IRI | Fantasizing | -0.004 | 0.032 | 0.500 | 0.291 | 0.410 |
|  | Perspective taking | -0.003 | 0.029 | 0.500 | 0.284 | 0.397 |
|  | Empathic Concern | 0.011 | 0.032 | 0.500 | 0.318 | 0.466 |
|  | Personal Distress | -9.326e -4 | 0.026 | 0.500 | 0.282 | 0.393 |
| MAS | Power-Prestige | -2.308e -4 | 0.027 | 0.500 | 0.289 | 0.406 |
|  | Retention-time | -0.010 | 0.032 | 0.500 | 0.310 | 0.450 |
|  | Distrust | 0.021 | 0.039 | 0.500 | 0.386 | 0.630 |
|  | Anxiety | -0.018 | 0.041 | 0.500 | 0.350 | 0.539 |

### 10. Results of the fMRI analysis for Outcome and PE_S_

**Supplementary Table 5.**
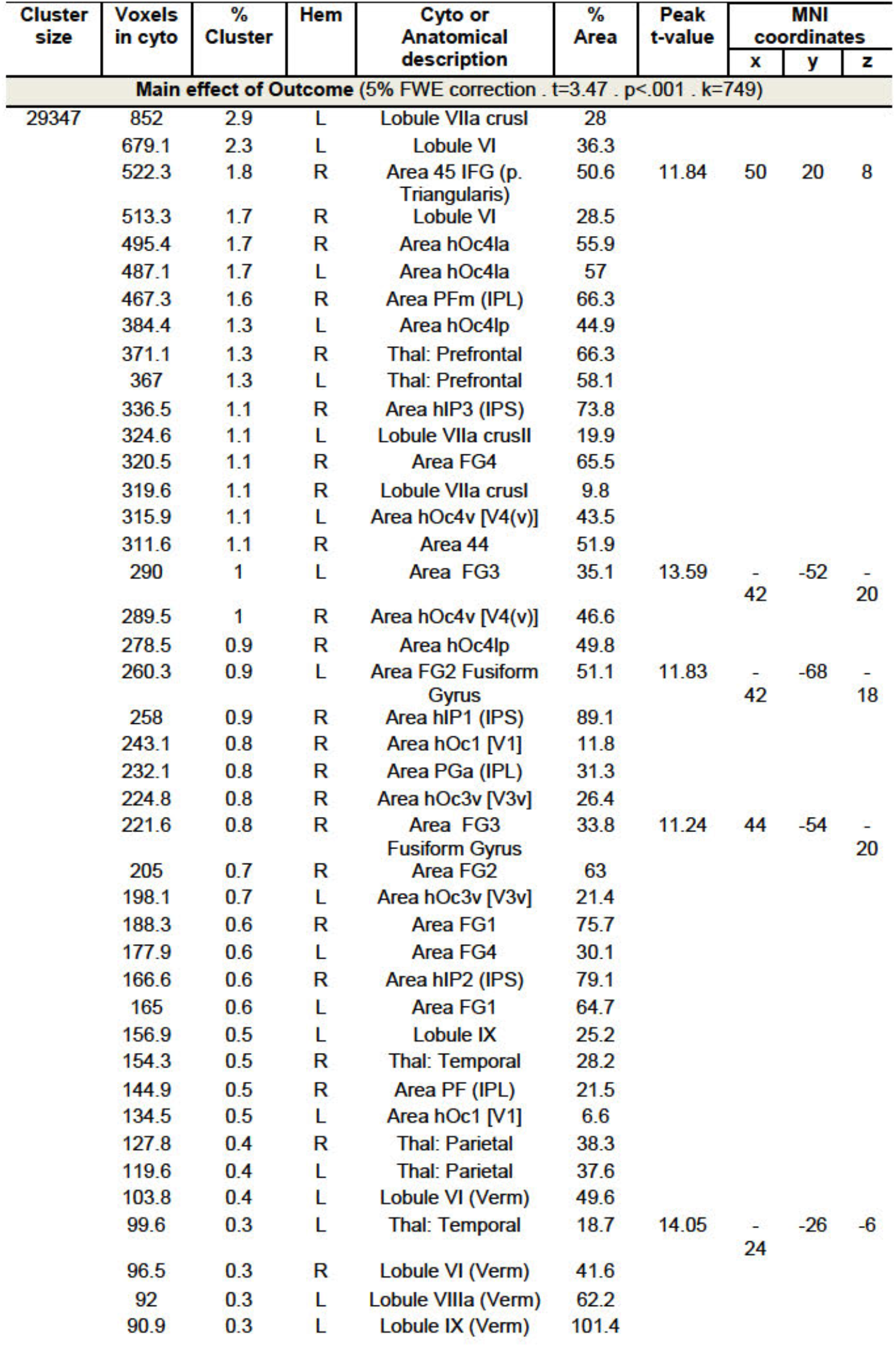
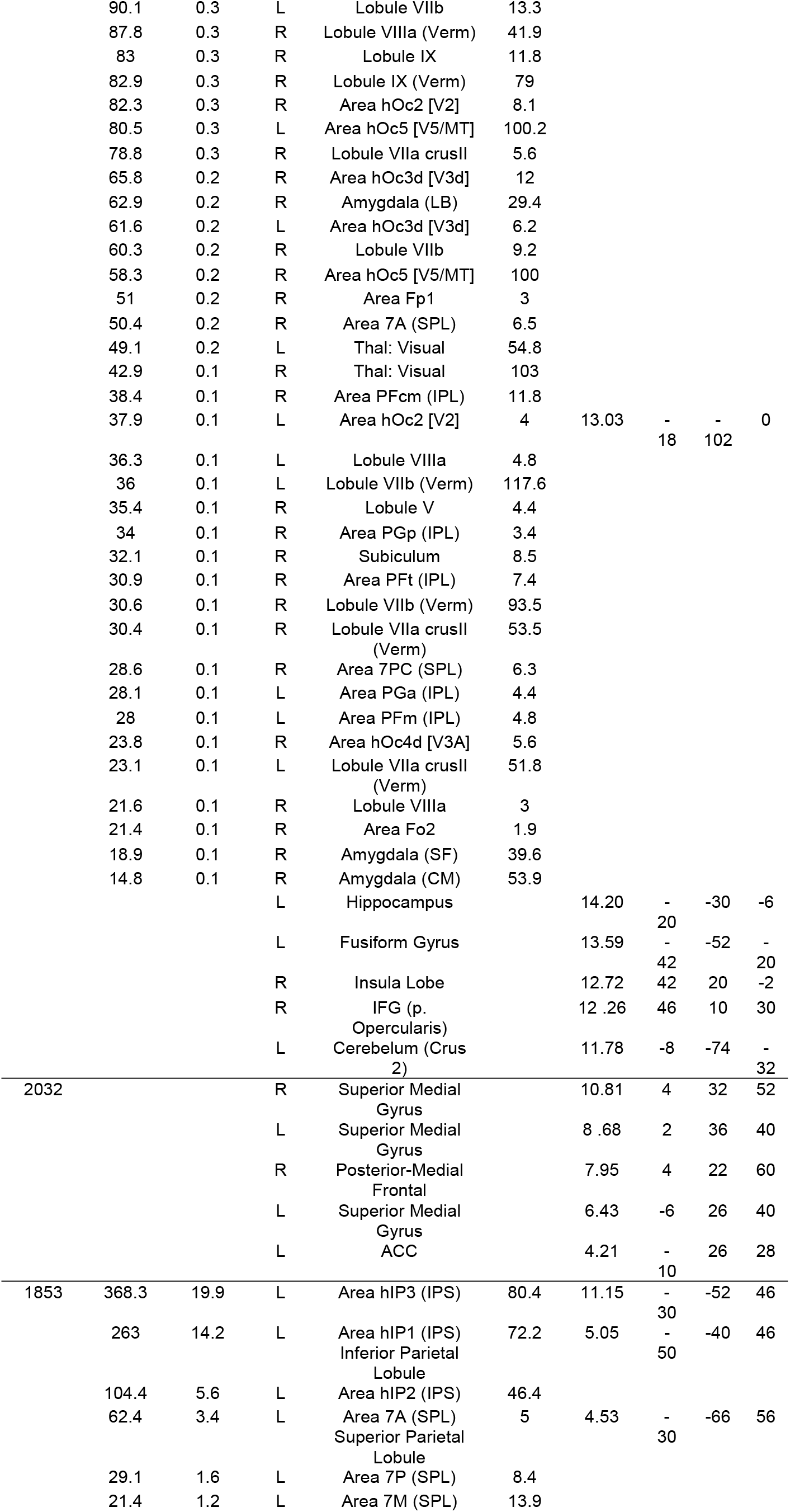
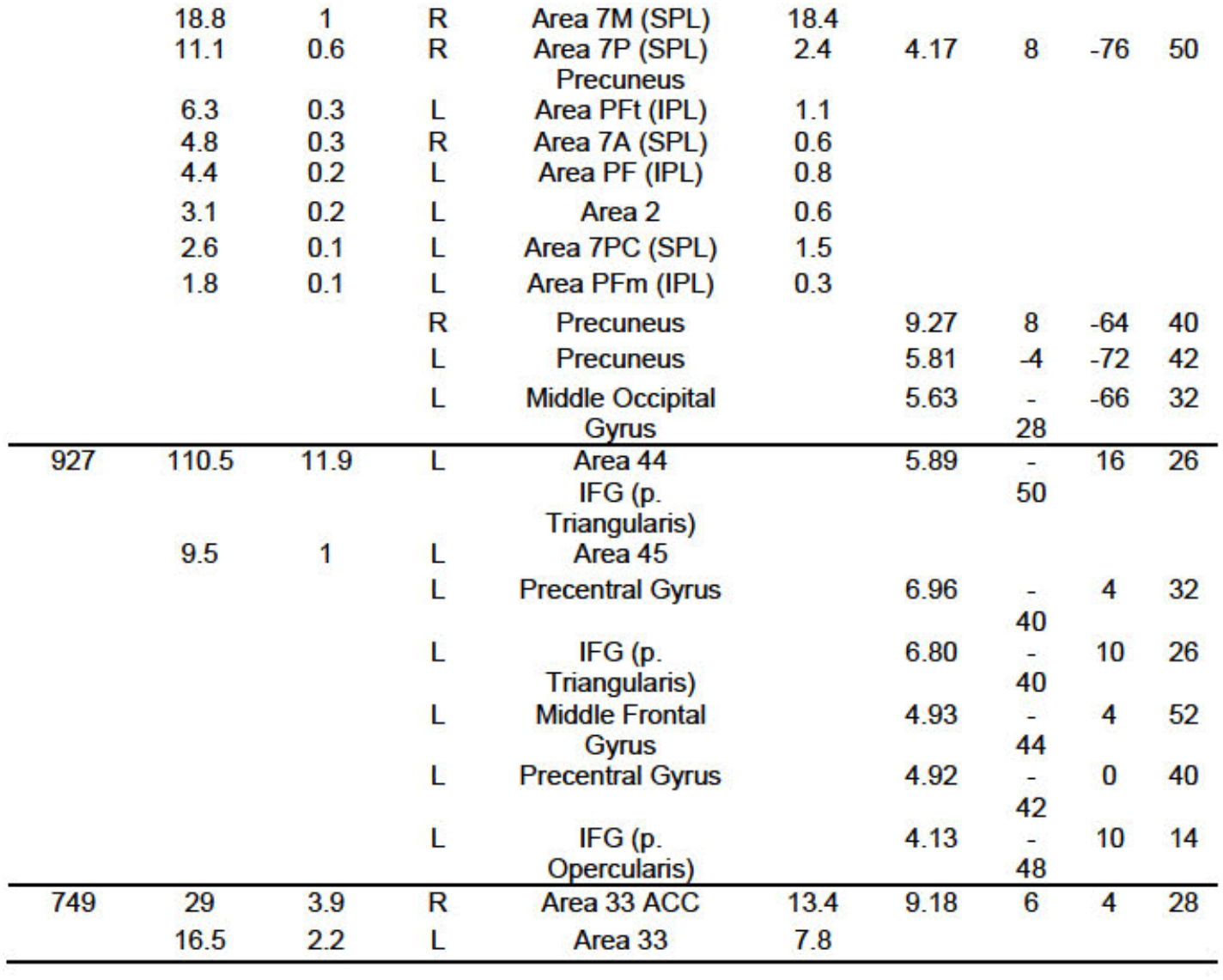
BOLD activity associated with the main effect of Outcome. Only clusters surviving a 5% FWE correction at the cluster size are reported (t=3.47, p < .001, cluster size 749; resampled voxel size: 2×2×2mm; Supplementary Figure 2). Brain regions are identified using the Anatomy Toolbox^30^. The columns refer to the size in voxels of each cluster; the number of voxels of that cluster falling within a specific cytoarcliitectonic region; the percentage of voxels in that region; hemisphere; cytoarchitectonic region (if available) or macro-anatomical description of the region; percentage of cytoarcliitectonic region activated by cluster; peak t-value within a particular region; and MNI coordinates of the peak. If more peaks were identified within the same cyto-architectonic or anatomical region only the peak with the highest t-value was included in the table. Peaks falling Outside the gray matter are not included in the table. Cyto architectonic description is only reported when a voxel has a probability over 40% to fall in that area and only for cyto-architectonic areas available in the anatomy toolbox; anatomical description is otherwise reported. The ‘*.txt’ file generated by the Anatomy toolbox can be found at: https://osf.io/rk8w4/?view_only=98b193a58aff48dda40b9d3d91ac5254

**Supplementary Table 6.**
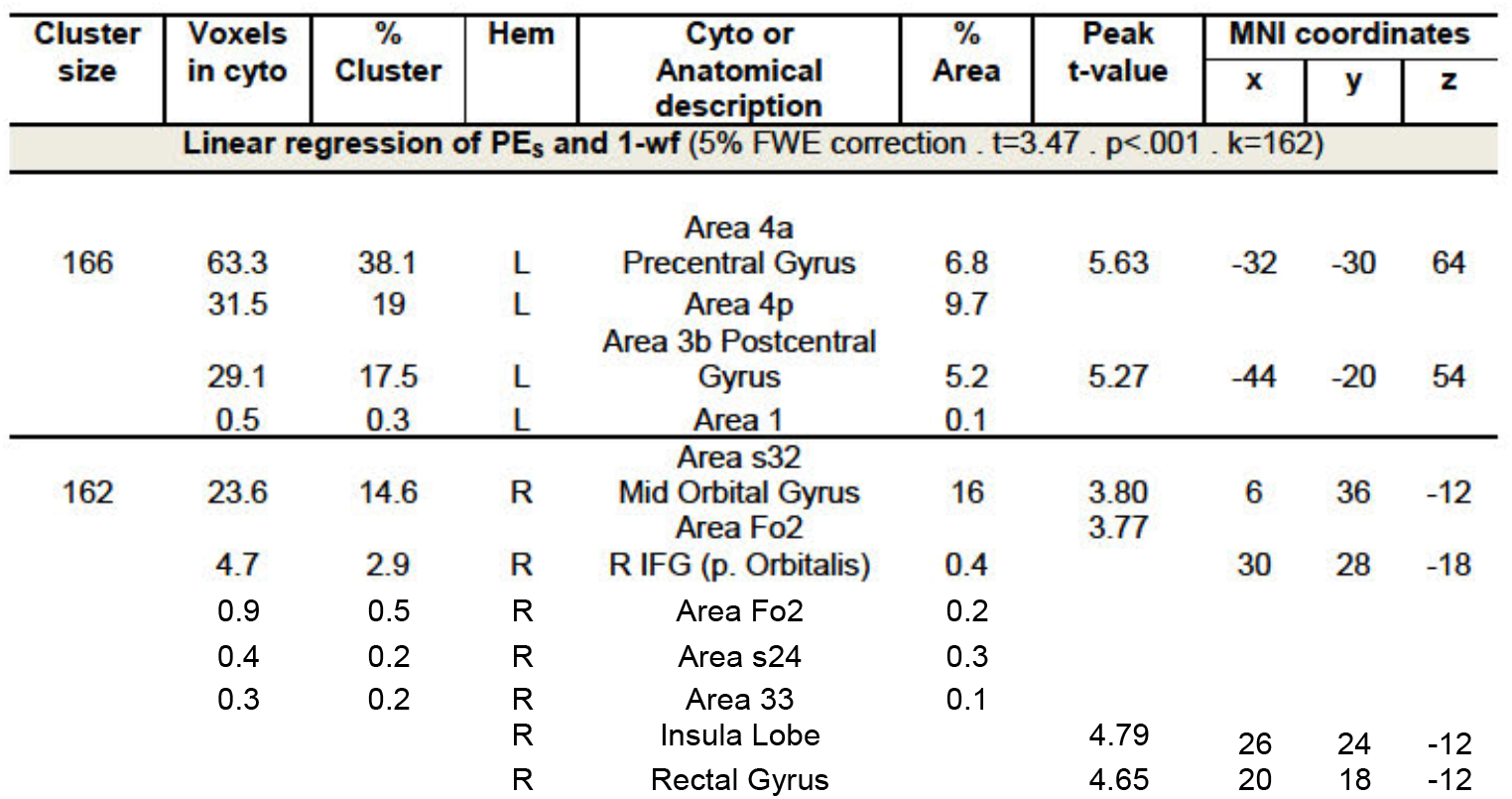
BOLD activity covarying with PE_S_ in a way that depends linearly on wf. Results of the linear regression of PE_S_ and 1-*wf*. Note that because the PES values from M2Out have been divided by 1-*wf* prior to entering them into the first level parametric regressor, the PES parameter estimates would no longer depend on 1-wf if PES signals were similarly strong across participants. Only clusters surviving a 5% FWE correction at the cluster size are reported (t=3.47, p < .001, cluster size 162; resampled voxel size: 2×2×2mm; Supplementary Figure 2). Brain regions are identified using the Anatomy Toolbox^30^. The columns refer to the size in voxels of each cluster; the number of voxels of that cluster falling within a specific cytoarchitectonic region; the percentage of voxels in that region; hemisphere; cytoarchitectonic region (if available) or macro-anatomical description of the region; percentage of cytoarchitectonic region activated by cluster; peak t-value within a particular region; and MNI coordinates of the peak. If more peaks were identified within the same cyto-architectonic or anatomical region only the peak with the highest t-value was included in the table. Peaks falling outside the gray matter are not included in the table. Cyto architectonic description is only reported when a voxel has a probability over 40% to fall in that area and only for cyto-architectonic areas available in the anatomy toolbox; anatomical description is otherwise reported. The ‘*.txt’ file generated by the Anatomy toolbox can be found at: https://osf.io/rk8w4/?view_only=98b193a58aff48dda40b9d3d91ac5254

**Supplementary Table 7.**
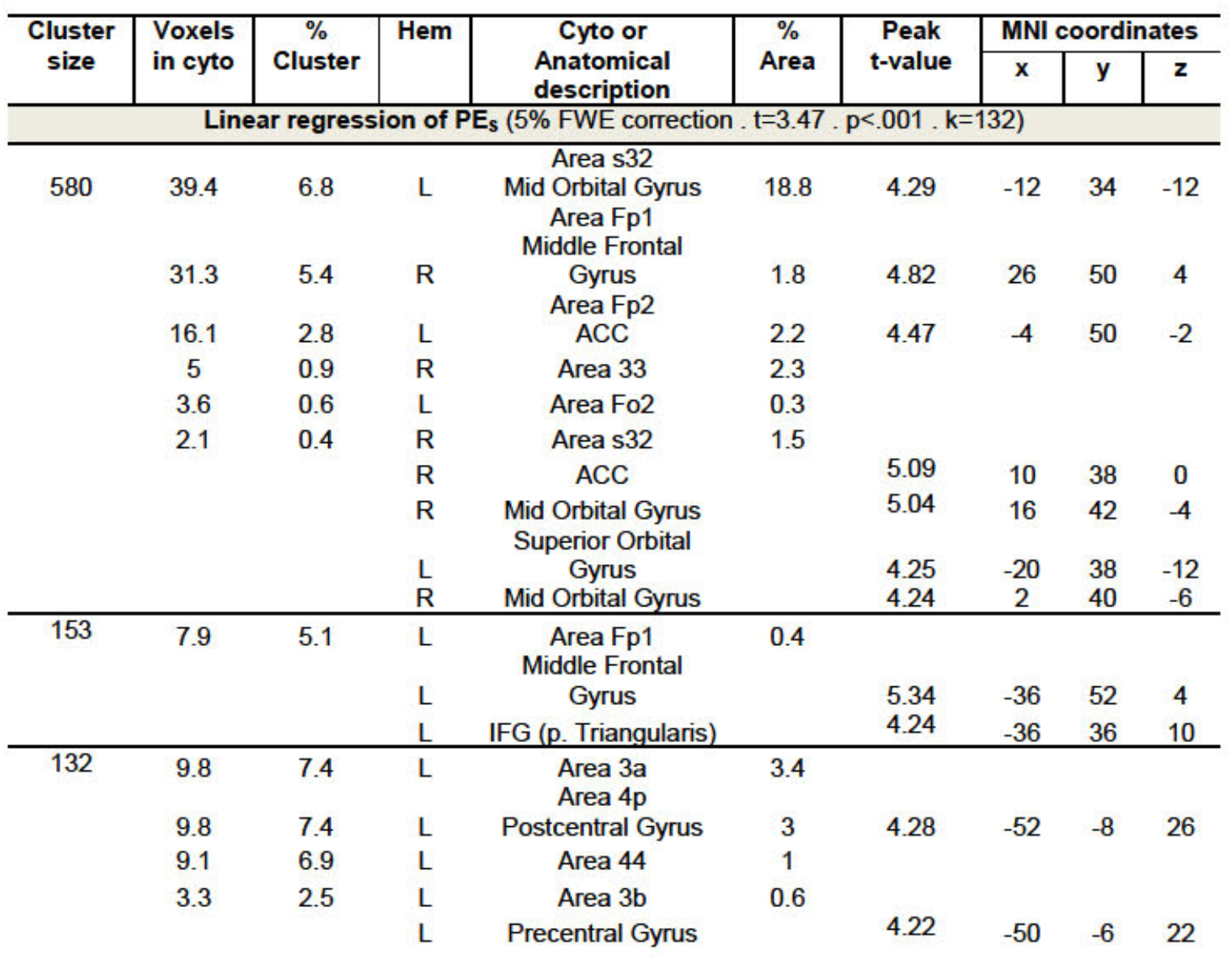
BOLD activity covarying positively with PES after removing variance explained by wf. Results of the constant in the linear regression of PE_S_ and 1-*wf*, capturing voxels where the parametric modulator for PES is above zero after removing variance explained by 1-*wf*. Note that because the PE_S_ values from M2Out have been divided by 1-wf prior to entering them into the first level parametric regressor, the PE_S_ parameter estimates used in this analysis would survive this contrast particularly well if all participants had similarly positive prediction errors. Only clusters surviving a 5% FWE collection at the cluster size are reported (t=3.47, p < .001, cluster size 132; resampled voxel size: 2×2×2mm; Supplementary Figure 2). Brain regions are identified using the Anatomy Toolbox^30^. The columns refer to the size in voxels of each cluster: the number of voxels of that cluster falling within a specific cytoarchitectonic region; the percentage of voxels in that region; hemisphere; cytoarchitectonic region (if available) or macro-anatomical description of the region; percentage of cytoarchitectonic region activated by cluster; peak t-value within a particular region; and MNI coordinates of the peak. If more peaks were identified within the same cyto-architectonic or anatomical region only the peak with the highest t-value was included in the table. Peaks falling outside the gray matter are not included in the table. Cyto architectonic description is only reported when a voxel has a probability over 40% to fall in that area and only for cyto-architectonic areas available in the anatomy toolbox; anatomical description is otherwise reported. The ‘*.txt’ file generated by the Anatomy toolbox can be found at: https://osf.io/rk8w4/?view_only=98b193a58aff48dda40b9d3d91ac5254

### 11. fMRI Parameter Recovery

The average correlation between the time courses of the parametric modulators for PES and PEM was −0.26, ranging from −0.49 to −0.03. Due to this correlation, we explored whether our experimental design and GLM approach can disentangle voxels that represent PES from those representing PEM, and whether they can differentiate voxels linearly dependent on *wf* from those that are not. Our GLM included, during the outcome period, a boxcar for the duration of the movie with two parametric modulators, one for PES and one for PEM. Both have been normalized by dividing them with 1-*wf* and wf respectively. This was done, as described in the Methods and Materials section of the main manuscript, to ensure that PES and PEM predictors become independent of preference and *wf* per se. When used in the GLM, the parameter estimates for these normalised PES values can then be compared across participants to identify if the brains of participants with higher weight on shocks (i.e. larger value for 1-*wf*) show larger signals for a given outcome than participants with lower weight on shocks. Using the original PES values would make that interpretation difficult, because they are already dependent on *wf*.

For this parameter recovery, we ran 1000 simulations. In each, we simulated 25 participants. For each participant, we used their own design matrices (the same used for the actual GLM first level analysis of the fMRI activity after convolution with the haemodynamic response function) to mix signals in each subject using three mixings (i) −1*PES+0*PEM+noise; (ii) 0*PES+1*PEM+noise, and (iii) −1*PES+1*PEM+noise. Noise was a random gaussian set at 1std of the mixed signal. Next, we ran a GLM using the same design matrix, and saved the parameter estimates for PES (we will call βPES) and PEM (we will call βPEM) for each participant. We then perform a t-test for βPES and one for βPEM to see if across the 25 parameter estimates (one per participant) there is evidence against the null hypothesis H_0_:βPES=0 or H_0_:βPEM=0. Of course, if PES was mixed into the voxels activity (case i or iii), a significant t-test would be a hit, while a non-significant t-test would be a miss, and the same applies to PEM for case ii. After repeating this procedure 1000 times, we count the proportion of the 1000 simulations where a t-test was significant against H_0_:βPES=0 or βPEM=0. Additionally, to see how often the analysis falsely detects a dependence on wf although wf was not included in the mixing, we also look at *r*(*wf*, βPES)=0 and *r*(*wf*, βPEM)=0. Initially, we use p<0.05 as a criterion, to look at the specificity and sensitivity for the case in which we explore responses in the AVPS, which is univariate. We also indicate proportions at p<0.001, but this time for a one-tailed test, in parenthesis, to provide results relevant for an explorative whole brain analysis where the cluster-cutting threshold was set at 0.001. The proportion of significant results was as follows:

**Supplementary Table 8:** Percentage of two-tailed t-tests significant at p<0.05 from the 1000 simulations using signals generated without multiplications with wf or (1-wf), and in brackets, the percentage of one tailed *p*_1-tailed_<0.001. The top row specifies how the signals were generated before adding 1sd of noise, the leftmost column, the null hypothesis that was tested in the hypothesis testing.

| **H<sub>0</sub>** | **mixing** | **-1·PES+0·PEM** | **0·PES+1·PEM** | **-1·PES+1·PEM** |
| --- | --- | --- | --- | --- |
| **βPES=0** |  | 99.8(92.4) | 4.4(0) | 99.8(90.6) |
| **βPEM=0** |  | 5.5(0.1) | 99.8(90.8) | 99.5(91.9) |
| **r(wf, βPES)=0** |  | 5.2(0.7) | 5.3(0.1) | 5.4(0.4) |
| **r(wf, βPEM)=0** |  | 4.5(0.2) | 5.1(0.1) | 3.8(0.0) |

We then repeated the same analysis, but this time multiplying the signals with (1-*wf*) and *wf* as indicated in Supplementary Table 9 to simulate cases of voxels where signal strength depends on preference.

**Supplementary Table 9:** Percentage of two-tailed t-tests significant at p<0.05 from the 1000 simulations using signals generated with multiplications with *wf* or (1-*wf*), and in brackets, the percentage of *p*_1-tailed_<0.001. The top row specifies how the signals were generated before adding 1sd of noise, the leftmost column, the null hypothesis that was tested in the hypothesis testing.

| **H<sub>0</sub>** | **mixing** | **-1·(1-wf)·PES+0·wf·PEM** | **0·(1-wf)·PES+1·wf·PEM** | **-1·(1-wf)·PES+1·wf·PEM** |
| --- | --- | --- | --- | --- |
| **βPES=0** |  | 99.3(81.1) | 3.5(0.3) | 99(80.5) |
| **βPEM=0** |  | 3.8(0.2) | 98.5(64.8) | 98.7(62.3) |
| **r(wf, βPES)=0** |  | 75.9(26.4) | 5.1(0.0) | 74.6(26.5) |
| **r(wf, βPEM)=0** |  | 5(0.0) | 98.3(77.3) | 98.5(79.3) |

The above tables explore evidence against the null hypothesis, but for univariate analysis we also ask whether we can actually provide evidence for voxels mixed without a certain factor that the GLM provides evidence for the null hypothesis using Bayesian statistics (8), using a bound of BF_10_<1/3. Using a Bayesian test, with n=25, we know that |t|<1 provides evidence in favour of H_0_: βPES=0 over H1: βPES≠0, and |r|<0.17 for H_0_: r(*wf* βPES)=0 over H1: r(*wf* βPES)≠0 (BF_10_<1/3, using default priors in JASP). We thus counted the proportion with evidence in favour of H_0_ over H1 in all cases using these bounds.

**Supplementary Table 10:** Percentage of BF_10_<1/3 from the 1000 simulations using signals generated without multiplications with wf or (1-wf). The top row specifies how the signals were generated before adding 1sd of noise, the leftmost column, the null hypothesis that was tested in the hypothesis testing.

| $H_0$ | mixing | -1·PES+0·PEM | 0·PES+1·PEM | -1·PES+1·PEM |
| --- | --- | --- | --- | --- |
| $\beta PES=0$ | | 0 | 69.6 | 0 |
| $\beta PEM=0$ | | 66 | 0 | 0 |
| $r(wf, \beta PES)=0$ | | 57.1 | 58.5 | 59.3 |
| $r(wf, \beta PEM)=0$ | | 58.0 | 56.8 | 58.4 |

**Supplementary Table 11:** Percentage of BF_10_<1/3 from the 1000 simulations using signals generated with multiplications with wf or (1-wf). The top row specifies how the signals were generated before adding 1sd of noise, the leftmost column, the null hypothesis that was tested in the hypothesis testing.

| $H_0$ | mixing | -1·(1-wf)·PES+0·wf·PEM | 0·(1-wf)·PES+1·wf·PEM | -1·(1-wf)·PES+1·wf·PEM |
| --- | --- | --- | --- | --- |
| $\beta PES=0$ | | 0 | 70.1 | 0 |
| $\beta PEM=0$ | | 68.1 | 0.1 | 0 |
| $r(wf, \beta PES)=0$ | | 2.0 | 57.5 | 3.6 |
| $r(wf, \beta PEM)=0$ | | 57.5 | 0.0 | 0.0 |

Summary: In our simulations, with 1sd of noise, we can detect voxels with signals linearly dependent on PES and/or PEM accurately: If we use alpha=0.05, as we would for the AVPS analysis, signals generated by including PES but not PEM are detected as representing PES in ~99% of cases, and only in ~5% of cases as representing PEM, and vice versa for voxels generated to include PEM but not PES. Using Bayesian statistics, we can even provide evidence in favour of the H_0_ for the former (βPES=0) in ~70% of cases, and the latter H_0_ (βPEM=0) in 66% of cases. Within our sample size, we thus have power to arrive at conclusions that match the way we generated the signals in the majority of simulations. Even when p<0.001 is used, as it would for our exploratory whole brain analysis, power remains decent.

With regard to linear dependence on wf, we find that for voxels generated with PES but not PEM signals, if the signal was generated using (1-*wf*) as a multiplier, a significant correlation is detected in 76% of cases, and when not used in the generation, a significant correlation is found in 5% of cases, while evidence for H_0_:r(*wf* βPES)=0 is found 57% of cases.

### 12. Psychological description of the difference between our learning models

All our learning models had in common that they learned by updating expected values for the two symbols using prediction errors, and additively combined self-money and other-shock. All of them captured individual variability in preference using a weighting factor (*wf*). What varied was when the two outcomes were combined. In our model-free version (M1), the outcomes themselves are combined into a single composite outcome value, which is already biased by the participants preference (Figure 2). Translated in psychological terms, this model would capture a model-free learning in which each symbol is associated with a value that captures how good or bad the outcomes for those symbols have felt in the past. In case of devaluation, this model-free learning doesn’t switch its preference to the symbol that has the highest expected value on the remaining quantity because it does not have an internal model that separates expectations for self-money and other-shock. In contrast, in our unbiased model-based version (M2Dec), participants track expectations separately for self-money and other-shock - independently of preference - and preference only plays out during the decision-phase. In psychological terms, this captures an unbiased model-based learning in which participants separately know how likely each symbol will lead to self-money or other-shock, respectively. Only when a decision must be taken, will they combine these predictions with the relative value that self-money and other-pain have for them personally, to come to a decision under conflict (Figure 2). In this version, the variable ‘expected value’ represents expectation in the objective units in which the outcome is coded, independently of whether this particular outcome is more or less valued by the participant in the conflict situation. Accordingly, in case of devaluation, the participant can base their decision on accurate predictions for the remaining quantity, and should have ~80% preference for the symbol with the highest expected value for the remaining quantity. Finally, in our *biased* model-based version (M2Out), participants also track separate expectations for both symbols in terms of self-money and other-shock, but they do so in ways that depend on their individual preference. Specifically, when self-money and other-shock outcomes are revealed, they are multiplied with their subjective weight (*wf* for self-money and *1-wf* for other-shock, Figure 2), and expectations are updated using these weighted values. At the decision-stage, the comparison between the two symbols is then done based on a simple sum of these already weighted expected values. Psychologically, this means that people have separate models for self-money and other-shock, but that the models predict the subjective value of each choice rather than the objective outcomes they are associated with. As a result, in case of devaluation, participants will have weaker preferences for the symbol that leads to more favorable outcomes if the outcome that remains was the outcome they valued less, than if the outcome that remained was the outcome they valued more.

### 13. Stress Tolerance short questionnaire (STSQ)

The Stress Tolerance Short Questionnaire is designed to measure the stress tolerance quotient and the participants’ anxiety level.

In the first part participants have to rate each item from 1 (always) to 5 (never), according to how much of the time the statement is true for them.

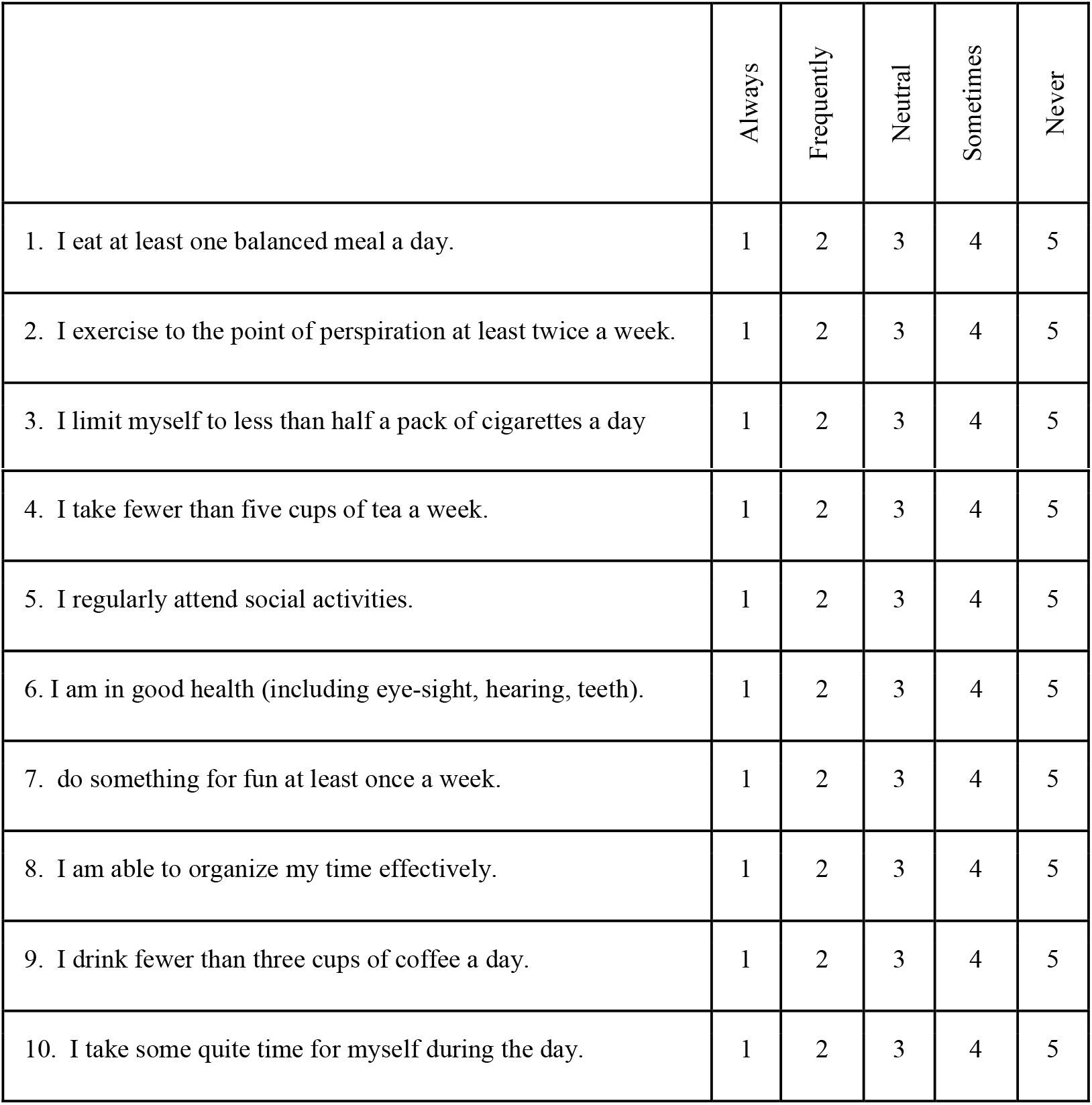

|  | Always | Frequently | Neutral | Sometimes | Never |
| --- | --- | --- | --- | --- | --- |
| 1. I eat at least one balanced meal a day. | 1 | 2 | 3 | 4 | 5 |
| 2. I exercise to the point of perspiration at least twice a week. | 1 | 2 | 3 | 4 | 5 |
| 3. I limit myself to less than half a pack of cigarettes a day | 1 | 2 | 3 | 4 | 5 |
| 4. I take fewer than five cups of tea a week. | 1 | 2 | 3 | 4 | 5 |
| 5. I regularly attend social activities. | 1 | 2 | 3 | 4 | 5 |
| 6. I am in good health (including eye-sight, hearing, teeth). | 1 | 2 | 3 | 4 | 5 |
| 7. do something for fun at least once a week. | 1 | 2 | 3 | 4 | 5 |
| 8. I am able to organize my time effectively. | 1 | 2 | 3 | 4 | 5 |
| 9. I drink fewer than three cups of coffee a day. | 1 | 2 | 3 | 4 | 5 |
| 10. I take some quite time for myself during the day. | 1 | 2 | 3 | 4 | 5 |

In the second part participants have to rate each item from 0 (not at all sure) to 3 (nearly every day), according to how often they have been bothered by the following problems over the last 2 weeks.

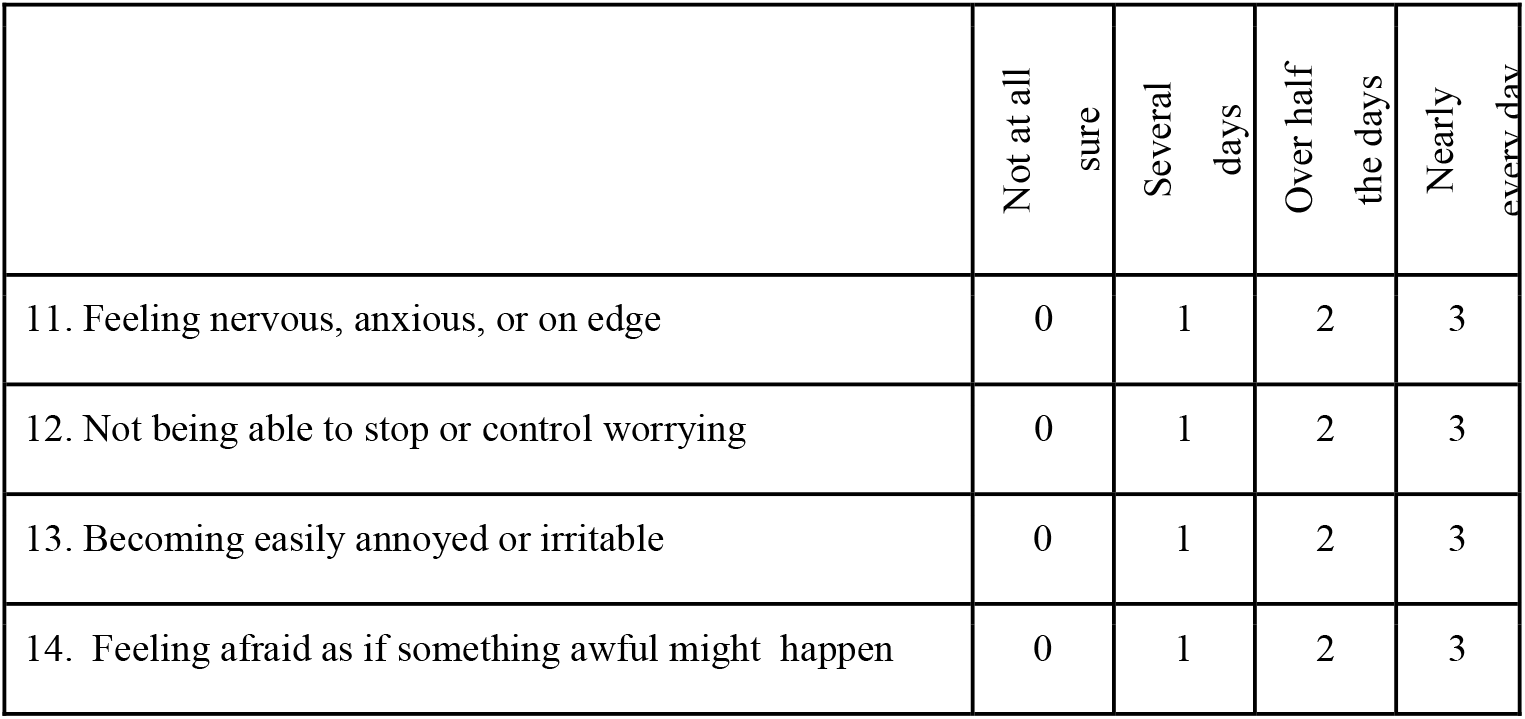

If participants obtained a score higher than 50 in the STSQ we would advise them not to continue with the experiment. This is because during the experiment they would be confronted with a number of possibly stressful videos, and put in a position in which their own benefit is detrimental to another individual. However, in case participants obtained a score higher than 50, we still gave them the possibility to continue with the task, after witnessing a sample video displaying a high intensity electrical stimulation to the confederate. None of our participants obtained a score higher than 50 in the STSQ.

### 14. Debriefing Questionnaire

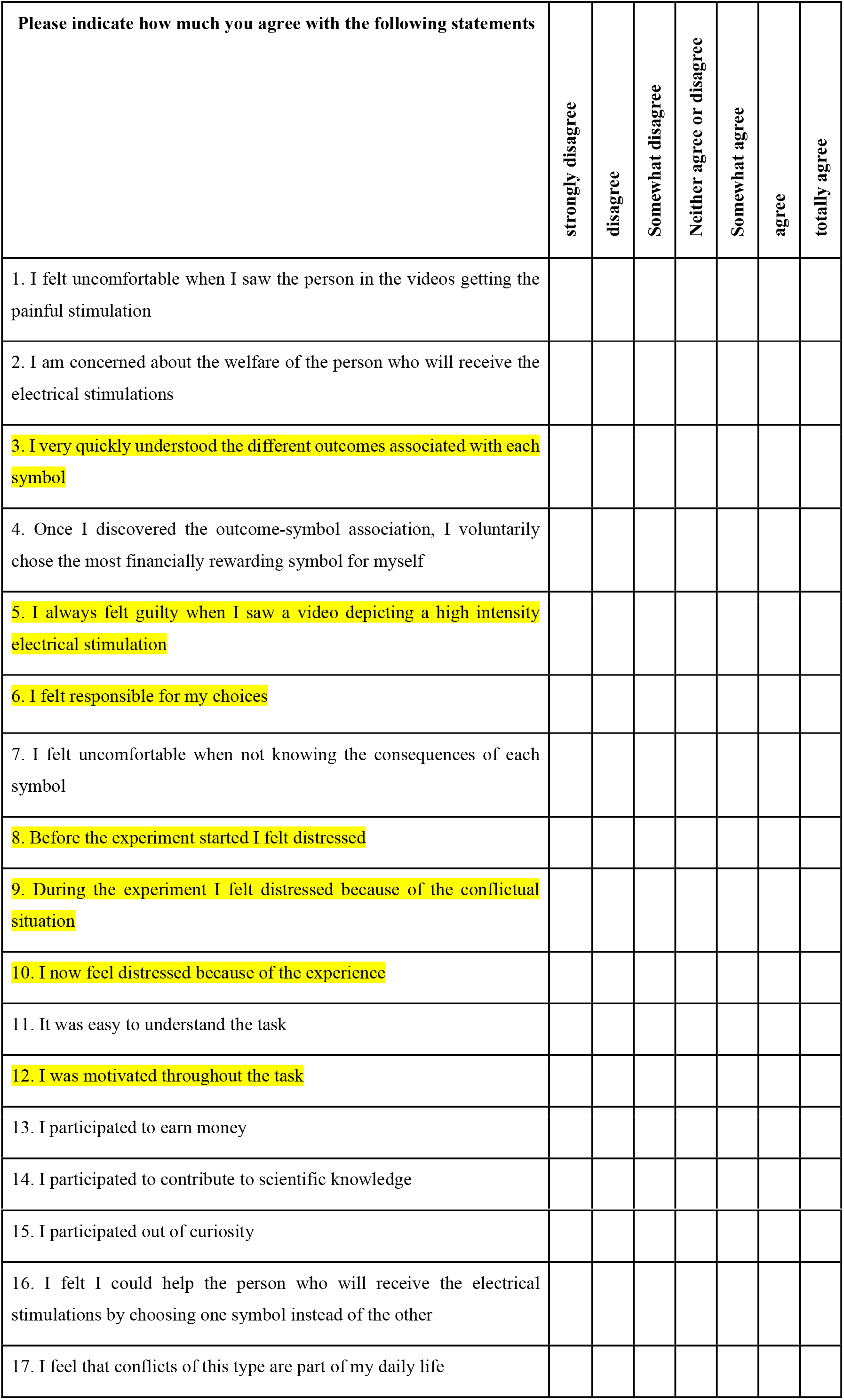

This questionnaire was administered after the learning task, to assess the motivation of the participants and the distress caused by the experiment. Highlighted in yellow are the questions which we considered more interesting to analyze, in relation to the behaviour in the learning task.

-Question 3: No effect of group on the declared understanding of the task (F_(2,76)_=0.53, p=0.594, BF_incl_=0.165).
-Question 5: effect of group on the sense of guilt when witnessing the high intensity electrical stimulations (Main effect of group on responsibility, F_(2,76)_=4.61, p=0.01, BF_incl_=3.77). Particularly Considerate declared to feel more guilty than both Lucrative (Independent sample t-test t_(51)_=2.85, p=0.01, BF_10_=6.93) and Ambiguous (Independent sample t-test t_(53)_=-2.38, p=0.02, BF_10_=2.44); while there was no evidence of difference between the amount of guilt reported by Lucrative and Ambiguous (Independent sample t-test t_(48)_=0.83, p=0.41, BF_10_=0.38).
-Question 6: absence of evidence of the effect of group on the amount of responsibility perceived during the task (Main effect of group on responsibility, F_(2,76)_=2.10, p=0.130, BF_incl_=0.564)
-Question 12: The three groups do not differ in the amount of motivation (F_(2,76)_=0.05, p=0.95, BF_incl_=0.115)
-Sum of the answers to question 8-9-10: No effect of group on the amount of stress caused by the task (F_(2,76)_=0.769, p=0.47, BF_incl_=0.201).

### 15. Eye gaze analysis

In an independent group of participants (N=41) we tested whether participants maximizing self-gain would look away from the facial expressions of pain (compared to participants minimizing the pain to others), to reduce their moral conflict. We collected eye gaze data through Prolific.sc. Specifically, we correlated the percentage of time spent by participants fixating the facial expression of pain videos in the Conflict condition, with the percent of considerate choices they made. As the percentage of fixation on the videos was not normally distributed (Shapiro-Wilk= 0.914, p=0.004) we calculated the Kendall’s ***τ***. Supplementary figure 7 shows the results of this correlation: despite a tendency in the expected direction, the relation was not significant and the Bayes factor leaned in favour of the null hypotheses. This preliminary data speaks against the idea that participants with more lucrative preference strongly avoided to look at the facial expressions of pain compared to participants with more considerate preference.

**Supplementary Figure 7.**
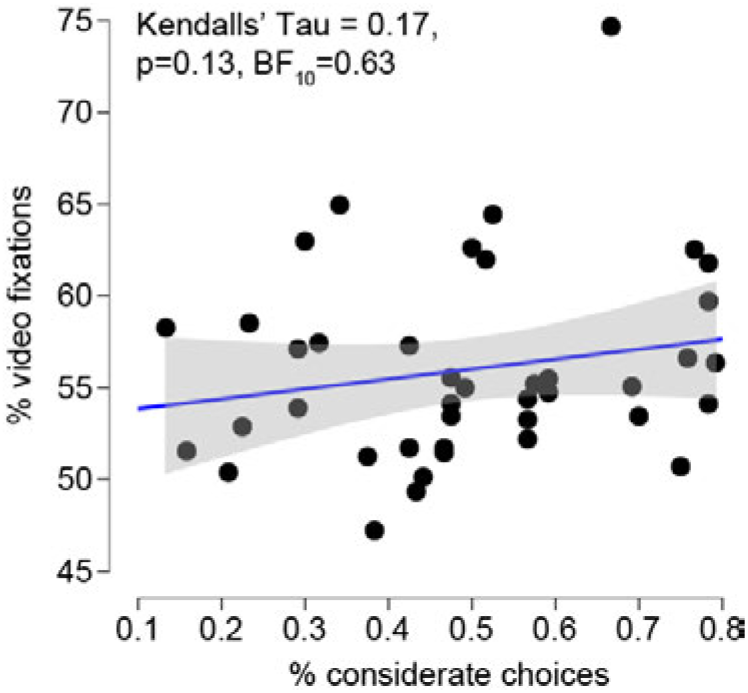
Correlation between time spent looking at the video and considerate choices in the Conflict condition.

---

**Figure 3_figure supplement 1.**
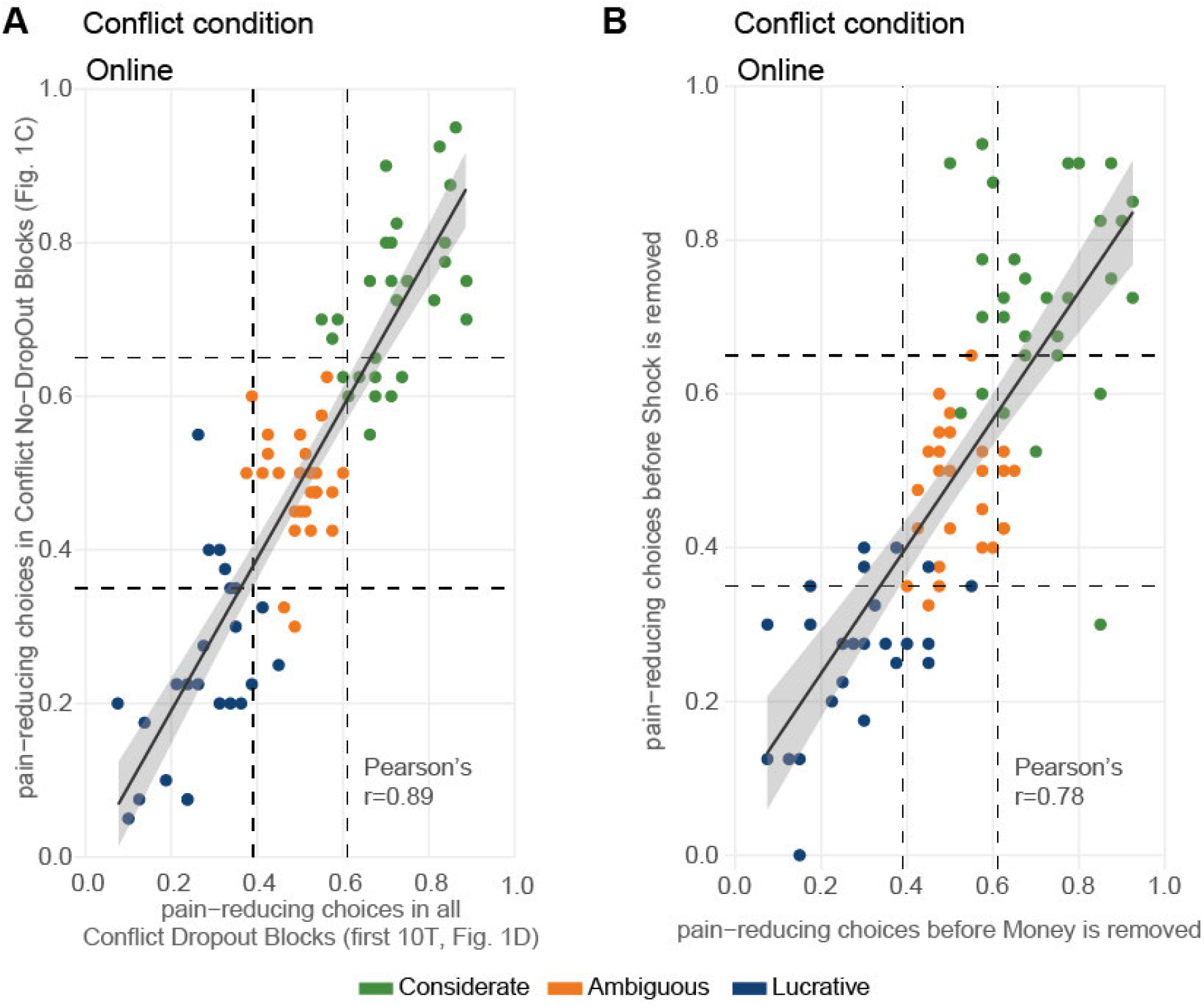
Choice allocation in DropOut and No-DropOut Conflict Online conditions. **(A)** Average choices allocation per participant during the first 10 trials of the 8 blocks that will later lead to DropOut (x-axis) and the 4 conflict No-DropOut blocks (y-axis). Pearson’s correlation= 0.89, t_(77)_=17.23, p<2.2e-16, BF_10_>1000; mean No-DropOut=0.501±0.025s.e m.; mean DropOut=0.512±0.023s.e m.; Paired sample test t_(78)_= 1.04, p=0.302; BF_10_=0.208. Color code represents the subgroup classification including all trials. The dashed lines represent the critical values for the binomial distribution *p*<0.05 separately for the DropOut and No-DropOut blocks. Regression line in black with shaded s.e.m. **(B)** As in (A) for the first 10 trials of the 4 DropOut blocks in which money will later be removed (x-axis), and the 4 DropOut blocks in which shock will later be removed (y-axis). Pearson’s correlation=0.78, p<0.001, BF_10_>1000; mean DropOut Money=0.52±0.024s.e m.; mean DropOut Shock=0.503±0.025s.e.m.; Paired sample test t_(78)_=1.21, p=0.23, BF_10_=0.25

**Figure 4_figure supplement 1.**
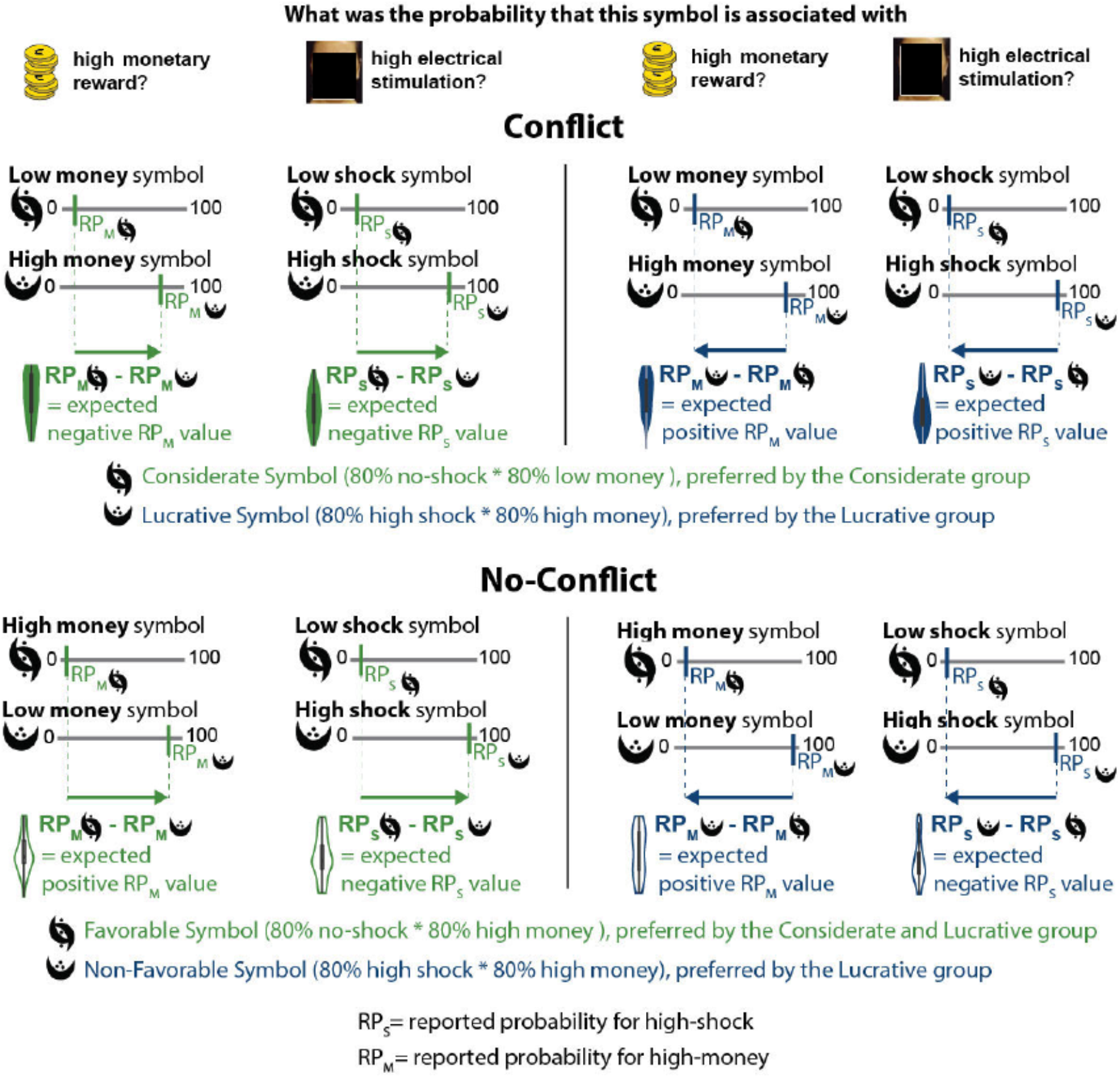
Reading key for Figure 4B. For each of the violin plots in Figure 4B, we here show how the violin plot is computed based on the difference between the two symbols.

